# When Prognostic Compatibility Does Not Guarantee Transfer Utility in Event-Limited Cross-Species Survival Modeling

**DOI:** 10.64898/2026.09.17.752483

**Authors:** Elena Krikun, Abedalrhman Alkhateeb

**Affiliations:** Department of Computer Science, Lakehead University, 955 Oliver Rd, Thunder Bay, ON P7B 5E1, Canada

**Keywords:** cross-species transfer learning, survival analysis, negative transfer, osteosarcoma, comparative oncology, domain adaptation, event scarcity

## Abstract

Cross-species molecular transfer may improve prognostic modeling in rare cancers, but under severe target-event scarcity the same data can easily be used to fit, choose, and evaluate adaptation, increasing the risk of negative transfer and model-selection bias. We evaluated canine-to-human osteosarcoma survival transfer using a chronologically frozen design. DOG^2^ served as the canine source domain, 50 MSigDB Hallmark modules defined the shared representation, and a known-truth benchmark comprised 180 scenarios and 21,600 replicates spanning 5–40 target events and multiple transport regimes. A frozen safety rule compared lowrank and module-selective adaptation; later diagnostics and a 6,000-replicate disjoint-seed mechanism experiment could not alter that decision. Neither selectable architecture met the negative-transfer limit (0.215 and 0.486 versus 0.10), although oracle analyses showed that the threshold was attainable. Prespecified post-HOLD weighting and threshold sensitivities showed that benchmark composition amplified the magnitude of A3 failure but did not create its instability, whereas A2 was comparatively insensitive to weighting. Training-set prognostic compatibility identified generator regimes (AUROC 0.980) but did not reliably order transfer benefit versus harm. Prediction-time gate hardening improved discrimination and reduced negative transfer, although its composite criterion was not met. Outcome-blind DOG^2^–TARGET analysis showed heterogeneous Hallmark preservation. In 86 TARGET cases with 29 events, the frozen canine classical model had IBS 0.159 versus 0.166 for a fold-local no-covariate Kaplan–Meier reference, while the human interpretation remained unresolved. Measurable prognostic compatibility therefore did not guarantee safe architecture-level transfer utility.

## 1. Introduction

Molecular prognostic modeling becomes difficult when the target disease is rare and the number of observed clinical events is small relative to the dimensionality of the molecular data. Osteosarcoma is a useful example of this problem. Public human cohorts with matched transcriptomic and survival data are limited in size, including the TARGET osteosarcoma cohort used in this study [1]. In such settings, the same small target dataset can easily be asked to fit a model, choose how much source information to borrow, tune the adaptation strategy, and evaluate the final predictor. This makes apparent improvements difficult to distinguish from model-selection effects.

Comparative oncology provides an additional source of molecular information. Osteosarcoma occurs spontaneously in pet dogs and shares molecular and microenvironmental features with the human disease. Cross-species genomic studies have identified conserved transcriptional programs and progression-associated signals in canine and human osteosarcoma [2, 3, 4]. More recent bulk and single-cell studies have also reported conserved tumor-microenvironment structure across species [5, 6].

These observations make canine osteosarcoma a biologically motivated source domain for studying molecular transfer to human disease.

Transfer learning is designed for settings in which labeled target-domain data are scarce but related source-domain information is available [7]. Related approaches have also been developed specifically for survival analysis, including transfer under the Cox model [8, 9]. Transfer is nevertheless not guaranteed to improve target prediction. If the source and target differ in the features or effects that matter for the task, source information can reduce target performance, a phenomenon commonly described as negative transfer [10, 11]. This risk is especially relevant when the target has only a small number of events, because flexible adaptation can itself be difficult to estimate and validate.

Molecular similarity between two domains does not resolve this problem by itself. A source representation can remain measurable in the target while the prognostic effects encoded by that representation are only partly transportable. Conversely, visible domain shift does not imply that all source information should be discarded. A useful transfer strategy therefore has to answer two distinct questions: whether source and target molecular structure is compatible, and whether borrowing source information actually improves target prediction relative to a target-only model. These quantities need not be the same.

Real cohorts alone cannot fully resolve this distinction. In a target cohort with only a few dozen survival events, observed prediction performance does not reveal which underlying molecular components are genuinely transportable, and the true source–target effect relationship is unknown. The mechanisms that can produce successful, neutral, or harmful transfer are therefore not directly observable from a single real dataset. We consequently used known-truth simulations as the primary environment for evaluating transfer safety and mechanism. In that setting, transportable fractions, effect concordance, sign reversal, covariance shift, mapping error, censoring, and event scarcity could be varied under controlled conditions while performance was evaluated on independent target outcomes. Real human data were then used at a separate evidence level rather than as the environment in which the transfer architecture was selected.

A second challenge is that negative-transfer protection is itself a model-selection problem. If a safety threshold or adaptation rule is changed after its performance is known, an unsuccessful architecture can be rescued by redefining the criterion. The opposite problem also matters: a negative result has limited meaning if the predefined safety requirement was impossible to satisfy within the benchmark. We therefore separated architecture selection, threshold-attainability analysis, diagnostic analyses, controlled mechanism experiments, and human evaluation into chronologically distinct stages. Rules for each stage were frozen before the quantities used at that stage were read, and analyses defined after an earlier result were explicitly assigned a weaker evidence status rather than being presented as part of the original architecture-selection test.

The study used the DOG^2^ canine osteosarcoma cohort as the source domain and an outcome-blind dog-to-human ortholog bridge represented by 50 MSigDB Hallmark gene sets. Candidate strategies included target-only survival learning, outcomezero-shot source transfer, target-head retargeting, low-rank adaptation, module-selective borrowing, and full neural finetuning. The primary known-truth benchmark comprised 180 scenarios spanning target event budgets from 5 to 40 events and multiple forms of source–target heterogeneity. Unlike selection by average discrimination alone, the frozen decision rule explicitly constrained negative and catastrophic transfer.

This study makes four methodological contributions. First, it provides a chronologically isolated evaluation protocol for cross-species survival transfer under event scarcity, with model definitions, safety criteria, and later diagnostic stages assigned explicit evidence levels. Second, it uses a known-truth benchmark to separate training-set prognostic compatibility from actual architecture-level transfer utility and to evaluate whether the frozen safety criterion is itself attainable. Third, it uses an independent set of simulation seeds for a controlled mechanism experiment in which predefined interventions test how head adaptation, source centering, selective borrowing, and prediction-time gate hardening affect transfer behavior. Fourth, it adds a secondary outcome-blind analysis of real canine and human expression structure, using all 50 Hallmark modules together with independently frozen biological anchors, without using human survival outcomes to define the biological comparison.

The resulting design addresses four questions. Can a transfer architecture meet a prespecified negative-transfer safety rule when target events are scarce? Does measurable training-set prognostic compatibility provide a reliable surrogate for the actual utility of transfer? When transfer behavior changes across source–target regimes, which aspects can be localized by controlled interventions without reopening architecture selection? Finally, how does the resulting computational setting relate to the biological organization of real canine and human osteosarcoma and to a human cohort evaluated under a frozen, descriptive protocol?

The emphasis is therefore not on obtaining a favorable result from a single small human cohort. Instead, the study asks under what conditions cross-species molecular borrowing can be evaluated as informative, unsafe, or indeterminate when target outcome information is limited. TARGET-OS is used as a descriptive human stress test rather than as a retrospective architecture-selection dataset, while the separate outcome-blind biological analysis characterizes the real cross-species expression setting without using TARGET survival outcomes.

## 2. Materials and Methods

### 2.1. Study chronology, prespecification, and outcome firewall

The study was organized as a sequence of frozen analysis stages so that later outcome information could not be used to revise earlier model definitions, selection rules, or interpretation thresholds. The chronology and evidence levels are summarized in Fig. 1 and Table 1.

**Table 1.** Evidence status of the main analysis layers.

| Analysis layer | Timing of specification | Outcome access | Permitted interpretation |
| --- | --- | --- | --- |
| Frozen simulation benchmark | Scenario grid, models, metrics, and selection rules fixed before simulation model fitting | No real human outcomes used | Primary evidence for negative-transfer safety and the frozen A2/A3 selection decision |
| Post-HOLD diagnostics | Each diagnostic scope fixed after HOLD and before its corresponding new diagnostic quantities were read | No real human outcomes used | Explanatory only; could not alter HOLD, change thresholds, or reopen model selection |
| New-seed H1–H4 experiment | Twelve cells, seven interventions, and H1–H4 fixed before fitting 6,000 new replicates | No real human outcomes used | Prospectively specified controlled evidence within the frozen generator family; not architecture rescue |
| Final sign-inversion and regime-shape audit | Rules fixed after H1–H4 were known but before the new audit quantities were read | No real human outcomes used | Bounded post-result mechanism audit; could refine interpretation but not change H1–H4 or reopen development |
| Biological-context analysis | Frozen after TARGET outcome opening but before inspection of the completed TARGET evaluation | DOG<sup>2</sup> and TARGET expression only; no TARGET survival values used | Secondary characterization of the cross-species biological/domain structure; not independent validation of TARGET performance |
| TARGET-OS evaluation | Models, preprocessing, resampling, metrics, and reporting rules frozen before endpoint opening | Primary TARGET OS endpoint only | Descriptive, non-confirmatory human stress test; no model selection |
| Post-opening KM null reference | Null-reference rule frozen after TARGET opening and before the new reference was calculated | Exact already-opened TARGET outer-training outcomes only | Descriptive scale reference for IBS; could not tune, select, or reclassify any model or branch |

**Figure 1.**
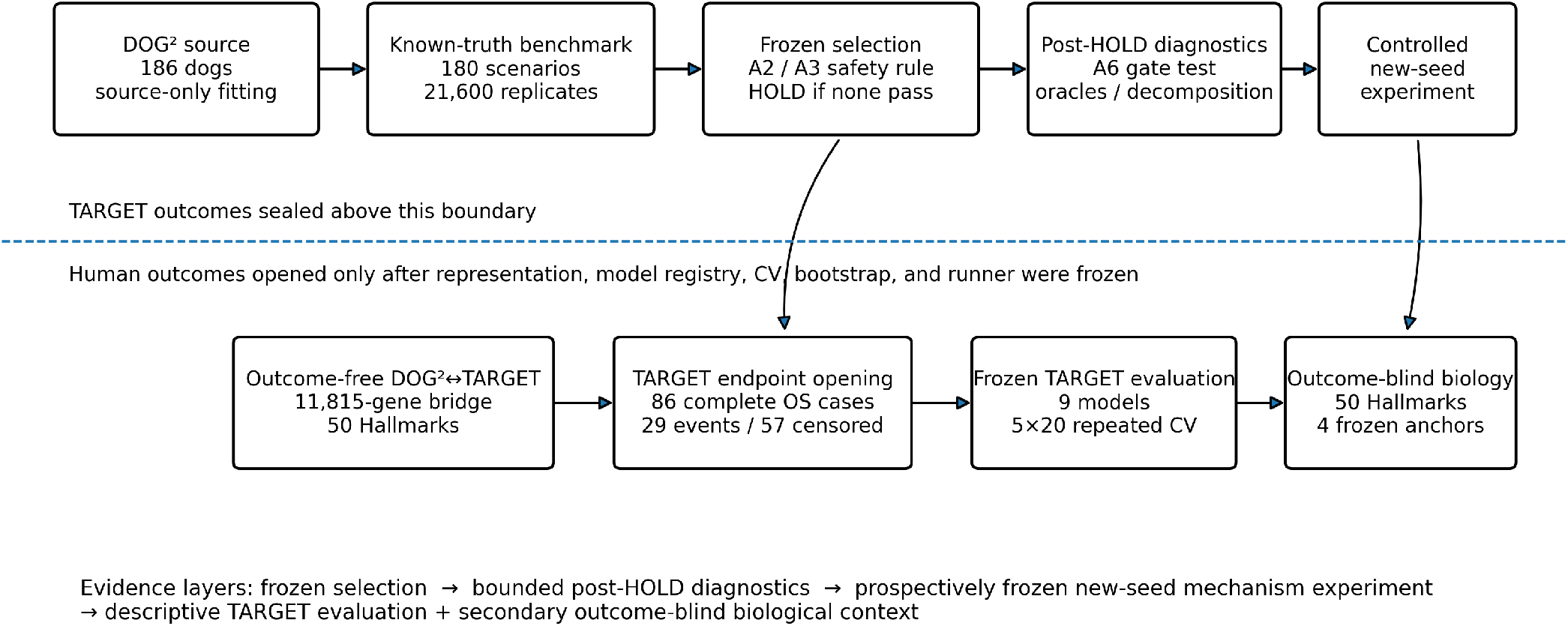
Chronology and outcome firewall of the cross-species transfer study. The known-truth simulation benchmark, frozen architecture-selection rule, post-HOLD diagnostics, and controlled new-seed mechanism experiment were completed before the descriptive human evaluation. The DOG^2^–TARGET expression representation, TARGET model registry, repeated cross-validation, bootstrap procedure, and executable evaluation runner were frozen before TARGET survival outcomes were opened. The subsequent biological-context analysis was outcome-blind and frozen before completed TARGET model results were inspected.

Cross-species feature construction and the initial model registry were defined without access to the reserved human outcome cohorts. GSE16091 was used only during a pre-TARGET premise stage to assess whether canine molecular information could be transported to a small human osteosarcoma cohort under a low-capacity classical model [2]. It was not used to rank neural architectures, choose simulation regimes, or tune the later transfer models. TARGET-OS, GSE21257, and GSE39055 outcomes remained closed during architecture development.

The primary architecture-selection environment was a known-truth simulation benchmark whose scenario grid, model registry, metrics, and selection rules were fixed before model fitting. The subsequent post-HOLD analyses were diagnostic: their purpose was to distinguish implementation problems, threshold attainability, and properties of the transfer setting, but they could not change the frozen architecture decision or reopen model development. A separate mechanism experiment was then defined on non-overlapping simulation seeds, with its cells, interventions, and H1–H4 support rules fixed before those new models were fitted.

A final sign-inversion and regime-shape audit was performed at a weaker evidence level. Its rules were fixed after H1– H4 were already known but before the corresponding sign-inversion and regime-shape quantities were inspected. This audit could refine the mechanism interpretation but could not change H1–H4 or create another simulation branch.

TARGET-OS was opened only after the expression representation, source models, resampling scheme, model registry, metrics, and evaluation runner had been frozen. The TARGET analysis was explicitly designated descriptive and nonconfirmatory; no architecture could be selected from it. The biological context analysis was frozen after TARGET outcome access but before inspection of the completed TARGET model results. It used expression values only and is therefore reported as a secondary, post-opening, pre-result-specified outcome-blind analysis. GSE21257 and GSE39055 outcomes remained sealed.

### 2.2. Cohorts, ortholog bridge, and Hallmark representation

The canine source cohort was the DOG^2^ osteosarcoma transcriptomic cohort (GEO GSE238110), containing tumors from 186 dogs with clinical follow-up [3]. The frozen source overall-survival endpoint contained 124 events. Human descriptive evaluation used the NCI TARGET osteosarcoma project [1]. The outcome-blind TARGET expression roster contained 88 cases. The frozen primary overall-survival analysis contained 86 cases with complete OS information, comprising 29 events and 57 censored observations. GSE16091 contained 34 human osteosarcoma samples with 15 OS events and was used only in the earlier premise stage [2].

Cross-species features were defined without outcome information. Canine transcript features were mapped to human genes using a versioned Ensembl orthology snapshot [12]. A feature was retained only when the canine identifier could be resolved deterministically, a unique dog-to-human one-to-one orthology was available, and the corresponding human gene was represented in TARGET. Ambiguous canine feature collisions, many-to-one orthologies, unresolved symbols, and duplicate human mappings were excluded by fixed outcome-blind rules. The resulting aligned expression universe contained 11,815 dog–human gene pairs.

Models used the 50 MSigDB Hallmark gene sets [13]. Within each training partition, gene means and population standard deviations were estimated from the training samples only. Genes with training standard deviation ≤ 10^−12^ were removed for that partition, and the fitted transformation was applied unchanged to held-out samples. For Hallmark module *m*, the score for patient *i* was

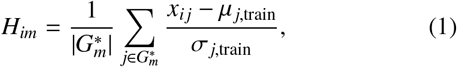

where *G*^*^_*m*_ is the subset of mapped genes that remained nonconstant in the training partition. At least 10 surviving genes were required for a module. Module-level means and standard deviations were subsequently fitted on the same training partition and applied unchanged to held-out samples.

Source and target domains were standardized independently. The frozen canine models were trained on Hallmark features standardized within DOG^2^, whereas TARGET models and outcome-zero-shot predictions used Hallmark features standardized from the current TARGET training partition. Thus, no whole-cohort TARGET statistics or held-out-patient information entered predictive preprocessing. The simulation engine used the same principle by standardizing source and target domains separately.

More aggressive domain-alignment procedures, such as correlation alignment (CORAL) [14], were not introduced after this representation had been frozen. A fold-local covariance-alignment method could in principle be constructed without using held-out outcomes, but it would constitute a different transfer transformation requiring its own prespecified implementation and evaluation. Independent within-domain standardization was therefore retained as the only cross-domain scaling step. More general cross-species representation methods can relax strict one-to-one gene matching [15], but adopting such a learned representation here would have changed the frozen feature space.

### 2.3. Transfer architectures and prognostic models

Survival models used the Cox proportional-hazards framework or a neural risk score optimized with the Cox partial likelihood [16, 17]. The frozen synthetic benchmark included a target-only classical model (B0), a classical residual-transfer comparator (B4), and five neural architectures (A0–A4). B4 regularized the target coefficient vector toward coefficients estimated from the canine source and served as a classical transfer comparator, conceptually related to coefficient-borrowing approaches for Cox-model transfer learning [9].

The neural source model used 50 Hallmark inputs, a 32-unit hidden layer, a 16-dimensional latent layer, hyperbolic-tangent activations, and a linear prognostic head. A0 trained the same architecture from scratch on the target. A1 froze the source encoder and re-estimated only the prognostic head. A2 froze the source network and added a rank-4 residual adapter in latent space. A3 used one continuous borrowing gate per Hallmark, allowing the source pathway to use the gated component and a target residual to model the complementary component. A4 initialized the complete neural network from the canine source model and fine-tuned it on the target.

For an A3 gate **g** ∈ [0, 1]^50^ and Hallmark vector **x**, the selective-borrowing risk score was

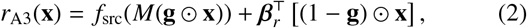

where *M* is the frozen cross-species mapping, *f*_src_ is the frozen source network, and *β*_*r*_ is the target residual coefficient vector.

A3 was evaluated only in the synthetic setting, where module transportability was known and an evolutionary borrowing prior could be defined before fitting. No corresponding per-module prior had been frozen for TARGET before human outcome access. A3 and its hardened version were therefore excluded from the real-data model registry rather than reconstructed after TARGET was opened.

Neural models were implemented in PyTorch [18] and optimized with Adam [19]. Architecture dimensions, learning rates, epoch counts, regularization strengths, and gradient-clipping rules were fixed before the simulation model matrix was fitted and were not tuned by scenario. No early stopping or scenario-specific hyperparameter search was used. The complete hyperparameter registry is provided in Supplementary Table S1.

The relationship between the synthetic architectures and the later TARGET comparators is summarized in Table 2.

**Table 2.**
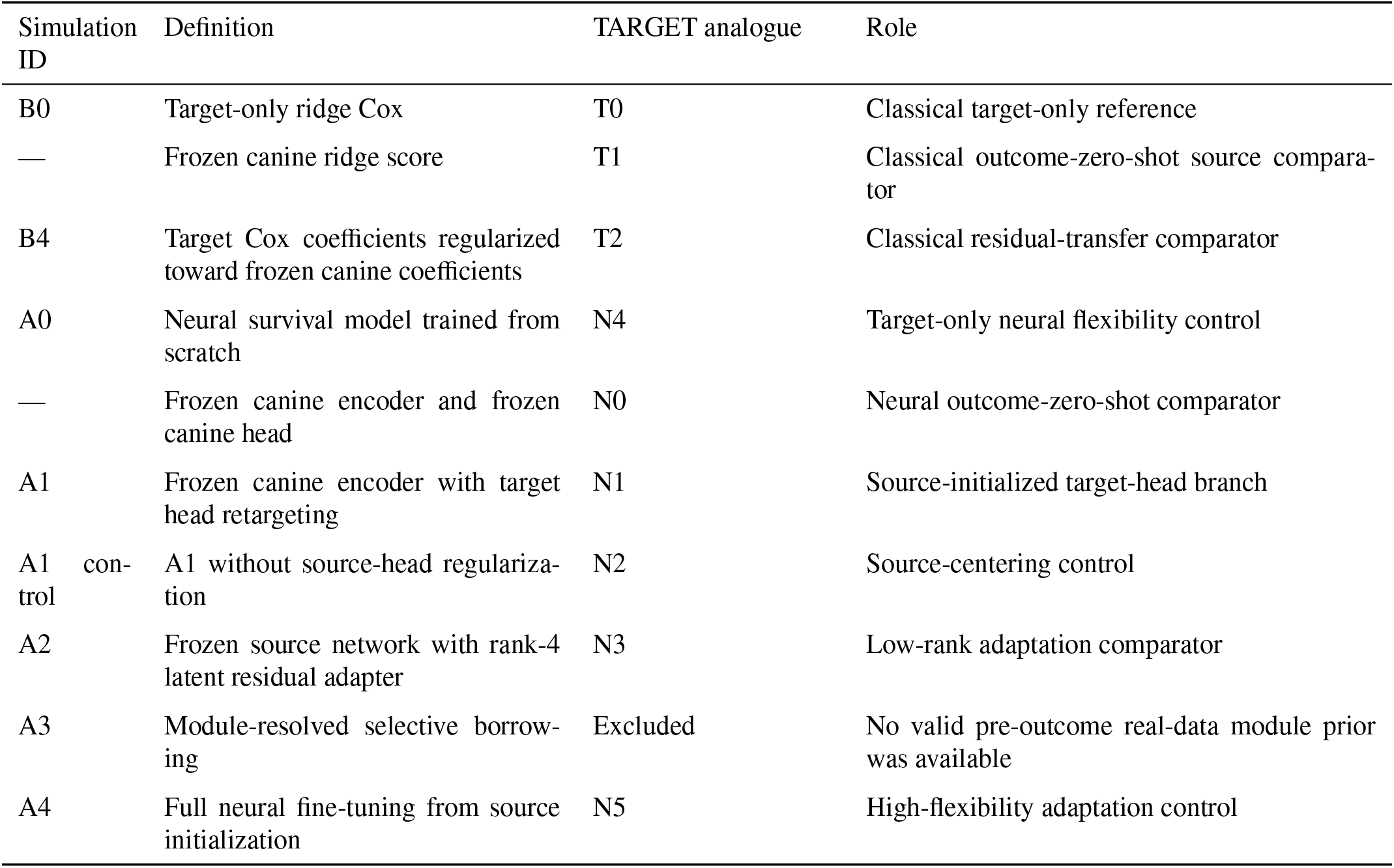
Frozen transfer-model families and their TARGET analogues.

| Simulation ID | Definition | TARGET analogue | Role |
| --- | --- | --- | --- |
| B0 | Target-only ridge Cox | T0 | Classical target-only reference |
| — | Frozen canine ridge score | T1 | Classical outcome-zero-shot source comparator |
| B4 | Target Cox coefficients regularized toward frozen canine coefficients | T2 | Classical residual-transfer comparator |
| A0 | Neural survival model trained from scratch | N4 | Target-only neural flexibility control |
| — | Frozen canine encoder and frozen canine head | N0 | Neural outcome-zero-shot comparator |
| A1 | Frozen canine encoder with target head retargeting | N1 | Source-initialized target-head branch |
| A1 control | A1 without source-head regularization | N2 | Source-centering control |
| A2 | Frozen source network with rank-4 latent residual adapter | N3 | Low-rank adaptation comparator |
| A3 | Module-resolved selective borrowing | Excluded | No valid pre-outcome real-data module prior was available |
| A4 | Full neural fine-tuning from source initialization | N5 | High-flexibility adaptation control |

### 2.4. Frozen simulation benchmark and architecture-selection rule

The simulation benchmark was designed to evaluate transfer under target-event scarcity before reserved human outcomes were used. Each replicate contained 50 modules, 10 of which carried source prognostic effects. The synthetic source cohort contained 186 observations and 124 events. Target training information was varied across event budgets of 5, 10, 15, 20, 29, and 40 events, and each replicate contained an independent target test population of 500 observations.

Six predefined source–target regimes ranged from fully transportable (R0), through intermediate levels of partial transportability (R1–R3), to nontransportable (R4) and misleading-source (R5) settings. The regimes varied the fraction of transportable causal modules, effect retention, and sign reversal. Additional axes varied covariance shift, censoring, cross-species mapping error, and correctness of the borrowing prior. Full generator parameters are reported in Supplementary Table S2. A legacy field in the released simulation artifacts labels R0–R3 as beneficial_regime; throughout the manuscript this label is interpreted only as a mechanistically compatible regime grouping and does not imply that transfer was empirically beneficial.

The complete benchmark contained 180 scenarios and 21,600 replicates. Core scenarios used 200 replicates and stress scenarios used 100. To prevent the different replicate counts from changing the model-selection target, metrics were first averaged within each scenario and then aggregated with equal weight across scenarios.

The primary discrimination metric was Uno’s concordance statistic for censored survival data [20], implemented with scikit-survival [21]. In the synthetic benchmark, the evaluation horizon was the 80th percentile of target-training event times, restricted to common train/test follow-up support. Secondary benchmark metrics included integrated Brier score (IBS) [22], one-dimensional Cox calibration slope, a horizon-specific calibration intercept (calibration-in-the-large), negative-transfer frequency, catastrophic negative-transfer frequency, and, for A3, recovery of the known module transportability mask. Calibration summaries were descriptive and did not enter the A2/A3 selection rule; their complete results are reported in Supplementary Table S3. Held-out risk-score standard deviation and percentile range were introduced only in later post-HOLD diagnostics and the controlled mechanism experiment.

For model *k*, negative transfer relative to B0 was defined by

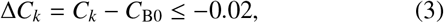

and catastrophic negative transfer by Δ*C*_*k*_ ≤ −0.05. A selectable candidate was required to have an equal-scenario negative-transfer rate no greater than 0.10 and a catastrophic rate no greater than 0.05 in both misleading-source and severe-shift settings. A3 additionally required mean transportability-recovery AUROC of at least 0.70 in partially transportable regimes and false borrowing no greater than 0.20 under misleading-source information.

Only A2 and A3 were eligible for final architecture selection. If both passed the protection criteria, Uno-C was compared first. An absolute Uno-C difference below 0.01 was treated as a practical tie, after which IBS was considered; an IBS difference below 0.01 resulted in selection of the simpler A2 model. If neither candidate met the protection criteria, the frozen rule selected no architecture.

The value +0.02 was not an original positive-utility requirement in this selection rule. It was introduced later as an operational reference for bounded post-HOLD diagnostics and was not used retrospectively to modify the original A2/A3 decision.

### 2.5. Post-HOLD diagnostic sequence and stop rule

A bounded post-HOLD sequence audited implementation integrity, threshold attainability, the relation between prognostic compatibility and held-out utility, and risk-score scale without refitting the original 21,600-replicate matrix, changing thresholds, or accessing reserved human outcomes. The training-set prognostic-compatibility statistic used a frozen source-derived risk score evaluated against the available target-training survival outcomes; it is therefore outcome-informed and is distinct from the later outcome-blind structural-concordance analysis of DOG^2^ and TARGET expression.

True-target-risk and regime-abstention oracles tested whether the frozen 0.10 negative-transfer requirement was attainable rather than relaxing it. A prospectively gated A6 trust/abstention branch was closed when its prespecified operational-compatibility AUROC criterion was not met. Comparator decomposition, regime-residualized and within-scenario associations, event-count abstention, and risk-scale/IBS diagnostics were explanatory only. Their exact chronology, stop rules, and pre-read/post-read boundaries are given in Supplementary Methods and Supplementary Table S4. A later post-HOLD strengthening contract added only descriptive robustness summaries. It exported the complete frozen B0/B4/A0–A4 family, recomputed A2/A3 under core-only and equal-regime aggregation while retaining the original 180- scenario estimand as primary, and evaluated neighboring harm definitions. For aggregation sensitivity, retained replicates were resampled within fixed scenarios to quantify Monte Carlo uncertainty; scenarios themselves were not resampled. The frozen −0.02 negative-transfer and −0.05 catastrophic-transfer definitions remained unchanged.

### 2.6. Controlled mechanism experiment on untouched simulation seeds

A focused mechanism experiment was specified after the original HOLD and diagnostic sequence but before any model was fitted on its new simulation replicates. Twelve cells crossed three target event budgets (10, 29, and 40) with four transfer regimes: R0, R2, R4, and R5. Covariance shift was fixed at zero, censoring at 0.40, mapping error at zero, and the borrowing prior was correct. Each cell contained 500 new replicates, giving 6,000 replicates in total. Their random seeds were verified to have no overlap with the original 21,600-replicate benchmark.

Seven interventions were frozen. M0 replayed the targetonly B0 model. M1 applied the frozen source neural network directly to the target without target fitting. M2 replayed A1. M3 was identical to M2 except that the penalty toward the source head was removed. M4 replayed the fitted A3 soft gate. M5 performed no additional fitting and replaced each fitted M4 gate only at prediction time by

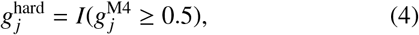

while retaining the already fitted residual coefficients. M6 used the true synthetic transportability mask as a binary gate and fitted only the complementary residual; it was an oracle and not an implementable model.

H1–H4 and their support rules were fixed before these 6,000 replicates were fitted. H1 compared M2 with M1 in misleading-source R5 and required an Uno-C gain of at least +0.02 at 29 events with a non-negative direction at 10 and 40 events. H2 compared M2 with M3 and required an Uno-C gain of at least +0.01 in the 29-event R5 cell, interpreted together with IBS and held-out risk-score standard deviation. H3 compared M5 with M4 and H4 compared M6 with M4; each required an Uno-C gain of at least +0.02 and an absolute reduction of at least 0.10 in catastrophic negative-transfer rate in the 29-event R5 cell. Exact contrast definitions are reproduced in Supplementary Table S5.

Uno-C was primary. Secondary quantities included IBS, held-out risk-score standard deviation, the 99th-to-1st percentile risk-score range, negative-transfer rate, and catastrophic negative-transfer rate. Uncertainty for the primary 29-event R5 comparisons was estimated using 5,000 paired replicate-level bootstrap draws.

After H1–H4 were known, a bounded arithmetic audit was specified to determine whether each component of the composite H3/H4 support rules was mathematically assessable given the realized control rate. This audit did not alter the original support thresholds or machine classifications.

A separate final audit was then specified to address sign inversion and regime shape. Its rules were frozen after H1–H4 were available but before the corresponding new audit quantities were read. M2 was compared with the sign-reversed score −*M*1. Sign-inversion equivalence required the complete paired 95% bootstrap interval to lie within [−0.02, +0.02]; evidence beyond sign inversion required a mean advantage of at least +0.02 with a positive lower confidence bound.

For the regime-shape audit, the 29-event M2 gain relative to B0 in R2 was compared separately with R0, R4, and R5. A discrete heterogeneity valley was defined only if all three bootstrap intervals for the other-regime-minus-R2 contrasts were above zero. This rule concerns four predefined generator categories and does not imply a continuous U-shaped biological relationship. No further simulation branch or diagnostic was permitted after this audit.

### 2.7. Secondary outcome-blind biological-context analysis

A secondary biological-context analysis was frozen after TARGET outcome opening but before inspection of the completed TARGET model results. It used expression values only and was explicitly classified as post-opening, pre-result-inspection, and outcome-blind. All 186 DOG^2^ and all 88 TARGET expression samples were used because no survival outcome or held-out prediction was involved.

For each of the 50 frozen Hallmark sets, Pearson gene–gene correlation matrices were computed separately in DOG^2^ and TARGET using the same ordered mapped genes. The primary structural-concordance statistic was Spearman correlation of the corresponding strict upper-triangle edge vectors,

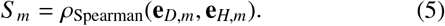

Each observed set was compared with 1,000 exact-size random panels matched on outcome-blind expression and variance ranks in both species. The complete 81-stratum matching algorithm, overlap audit, and fail-closed construction are reported in Supplementary Methods. Matched-control percentiles are descriptive, not *p*-values, and the overlapping Hallmark sets were not treated as 50 independent observations.

Four additional programs (M34, M40, M11, and M24) were carried forward as pre-existing coherence anchors from an earlier outcome-blind cross-cohort preservation analysis by the same authors. Their memberships and preservation labels had been finalized before the present biological-context analysis and were not selected from the current Hallmark results. M34 and M40 carried pre-existing labels of strong external canine representation preservation, whereas M11 and M24 carried noclear-preservation labels.

The anchor definitions were SHA-256-bound before the completed TARGET model-result summary existed. The biological-context contract also recorded a contemporaneous filesystem snapshot of the still-running TARGET evaluation, providing an auditable chronology for both the anchor definitions and the current structural analysis. Program sizes, prior labels, hash provenance, and the chronology evidence are reported in Supplementary Table S13 and the released provenance ledger. The anchors were evaluated with the same structural statistic and matched-control procedure and could not be removed or replaced if the new statistic disagreed with their prior labels.

This analysis characterizes outcome-blind structural concordance of the real cross-species expression setting; it is not independent evidence that such structure predicts prognostic transfer utility. A TARGET-derived module-level diagnostic was permitted only if an already frozen output admitted exact Hallmark decomposition without refitting or post-hoc attribution; otherwise it was omitted.

### 2.8. Descriptive TARGET-OS evaluation

TARGET-OS was a descriptive, non-confirmatory human stress test using the NCI TARGET osteosarcoma cohort [1]. Overall survival was the frozen primary endpoint. Of 88 expression cases, 86 had complete OS information (29 events, 57 censored); the single censored time-zero observation was retained unchanged after a bounded technical audit. Detailed endpoint reconciliation and the byte-level runner amendment are reported in Supplementary Methods.

All nine frozen models, T0–T2 and N0–N5 (Table 2), used the same event-stratified 5-fold outer cross-validation repeated 20 times. T0 and T2 used inner event-stratified tuning confined to outer-training patients; neural hyperparameters were not tuned on TARGET. Throughout this analysis, “outcome-zeroshot” means that no TARGET survival outcome was used to fit the source predictor; fold-local TARGET expression means and standard deviations remained part of the frozen representation and were estimated from the current outer-training partition only. T1 and N0 are outcome-zero-shot in this sense. Real-data A3 was excluded because no valid per-module prior had been frozen before human outcome access.

The primary metric was Uno-C [20], using the frozen 90th percentile of outer-training event times for time support. Secondary metrics were IBS and held-out risk-score scale. For risk models, the IBS calculation used an outer-training Breslow baseline [23] without refitting the risk coefficient, ranking, or calibration slope. Within each repeat, fold metrics were combined by sample-count weights and valid repeat estimates were then averaged; at least 16 of 20 repeats were required.

A later post-opening null reference contextualized IBS without changing any TARGET result. For each already-frozen outer partition, a no-covariate Kaplan–Meier curve was estimated from outer-training outcomes only and applied unchanged to every corresponding test patient. The original time grid, censoring/IPCW implementation, folds, and aggregation were reused exactly. The reference was descriptive only and could not affect model or branch selection.

Uno-C uncertainty and paired contrasts used 5,000 event-stratified patient-clustered bootstrap draws propagated through the fixed repeated-CV predictions; models and splits were not regenerated. Four prespecified reporting states (T-A–T-D) summarized the neural N0/N1 mechanism arm. If the N0 interval included 0.50 or a branch-defining paired interval spanned both −0.02 and +0.02, the result was mechanically T-D. The full model definitions, bootstrap mechanics, and branch rules are given in Supplementary Methods. No multiplicity adjustment was used because the complete TARGET arm was descriptive and no model-wise significance result could select a model.

GSE21257 and GSE39055 outcomes remained sealed and were not opened to explain, rescue, or reinterpret TARGET.

### 2.9. Reproducibility and audit trail

The analysis was implemented in Python using NumPy, pandas, PyTorch [18], and scikit-survival [21]. Simulation model fitting used GPU execution where defined by the frozen implementation; the TARGET evaluation used the frozen CPU execution path.

Random seeds, scenario definitions, model hyperparameters, cross-validation rules, interpretation thresholds, and output schemas were stored in versioned machine-readable artifacts. Scientific contracts and derived artifacts were bound by SHA-256 hashes, which were checked before downstream execution. Technical corrections made after an analysis boundary were implemented as new versioned amendments rather than by silently replacing the original artifact.

The Supplementary Material contains the frozen hyperparameter registry (Table S1), complete simulation-generator specification (Table S2), secondary simulation metrics including calibration summaries (Table S3), the full post-HOLD diagnostic sequence (Table S4), exact H1–H4 definitions and support rules (Table S5), biological matching and Hallmark-overlap audits (Table S6), TARGET fold/repeat-level metric and bootstrap details (Table S7), and the software, chronology, and artifact-hash ledger (Table S8). Byte-level provenance for the TARGET endpoint amendment is included in the Supplementary Methods.

Code, frozen configuration files, and the audit artifacts required to reproduce the reported analyses will be released with the final manuscript.

## 3. Results

The first five Results subsections report analyses from a single known-truth simulation generator family. The controlled mechanism experiment used new simulation seeds but retained the same generator family. The final two subsections relate these findings separately to real cross-species molecular structure and to the descriptive human outcome evaluation.

### 3.1. No prespecified architecture satisfied the frozen safety rule

Neither of the two architectures eligible for selection satisfied the frozen negative-transfer protection criteria across the 180-scenario benchmark. In the frozen candidate summary, the aggregate negative-transfer rate under equal-scenario weighting was 0.215 for the low-rank adapter A2 and 0.486 for the module-selective A3 model, compared with the prespecified maximum of 0.10. A2 also exceeded the catastrophic-transfer limit of 0.05 in both misleading-source scenarios (0.060) and severe-shift scenarios (0.078). The corresponding catastrophic-transfer rates for A3 were 0.712 and 0.353.

A3 nevertheless recovered the known transportable-module mask with a mean AUROC of 0.768 in partially transportable regimes and had a false-borrow rate of 0.186 under misleading-source information. Both values satisfied the frozen A3-specific criteria of AUROC ≥0.70 and false borrowing ≤0.20. Its safety failure therefore occurred despite meeting the prespecified module-recovery requirements. The frozen architecture rule selected no model and returned HOLD_NO_SAFE_SELECTABLE_ARCHITECTURE.

A negative selection result would be difficult to interpret if the safety criterion itself were unattainable. Post-HOLD oracle diagnostics were therefore used to assess attainability without changing the frozen threshold. A policy using the true target risk to determine whether to transfer had an aggregate negative-transfer rate of approximately 0.0001 and no catastrophic transfer. A regime-aware abstention oracle applied to A3 had a negative-transfer rate of 0.065 and a catastrophic-transfer rate of 0.009. Thus, the frozen 0.10 negative-transfer threshold was achievable within the simulated environment, although it was not met by either selectable architecture (Fig. 2).

**Figure 2.**
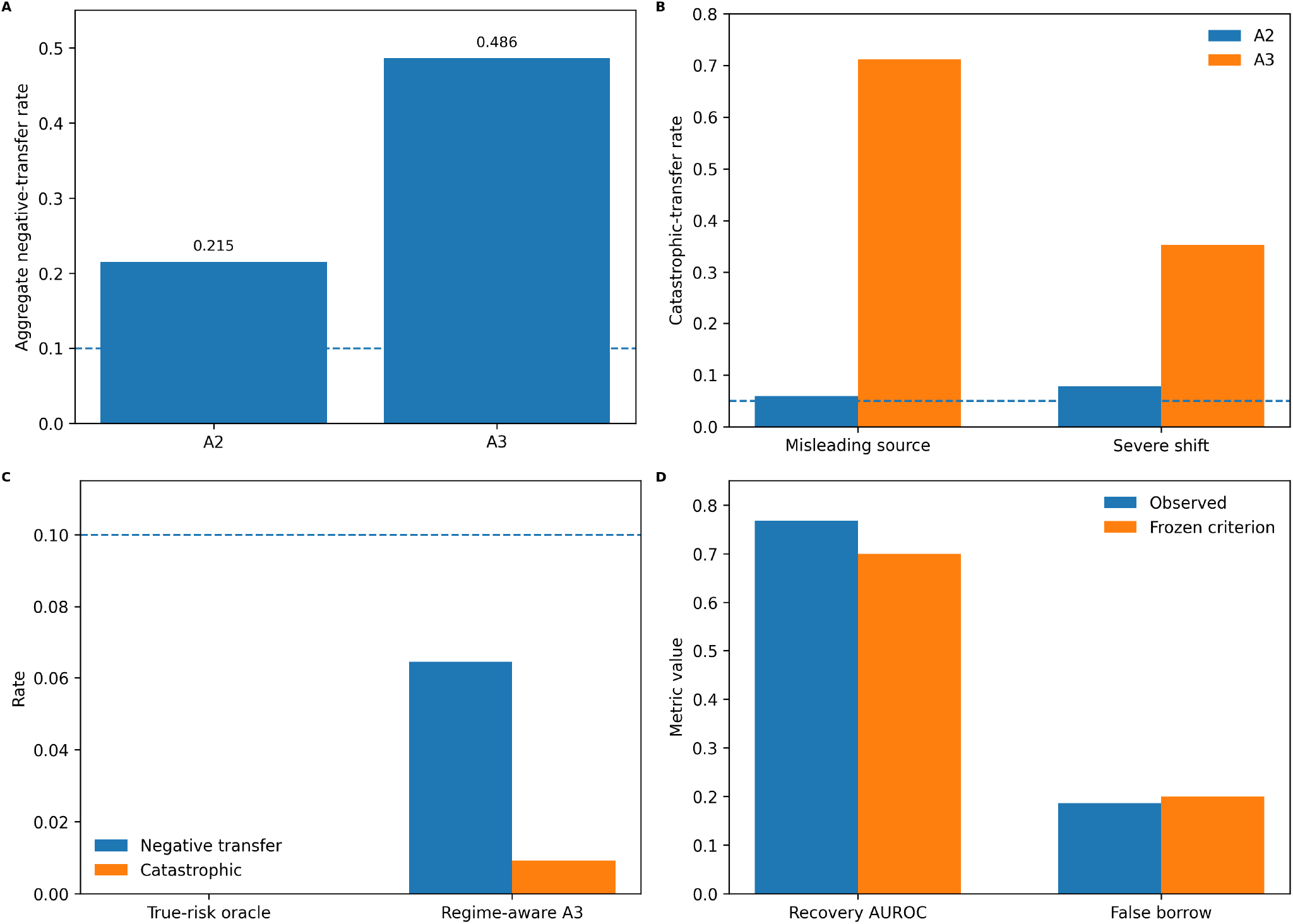
Frozen architecture-selection outcome and post-HOLD assessment of threshold attainability. **A**, Aggregate negative-transfer rates for A2 and A3 relative to the frozen 0.10 safety threshold. **B**, Catastrophic-transfer rates in misleading-source and severe-shift settings relative to the frozen 0.05 limit. **C**, Post-HOLD true-target-risk and regime-aware oracle references showing that the negative-transfer threshold was attainable within the known-truth benchmark. **D**, A3 module-recovery AUROC and false-borrow rate relative to their prespecified criteria.

The complete frozen seven-model family also showed that average discrimination and transfer safety were not aligned. A1 had the highest equal-scenario mean Uno-C (0.572) but an aggregate negative-transfer rate of 0.287, whereas the neural target-only control A0 closely matched B0 in mean Uno-C (0.555 versus 0.556) and had a negative-transfer rate of only 0.009. The full descriptive family summary is reported in Supplementary Table S9; nonselectable models remained nonselectable.

The post-HOLD aggregation sensitivity further distinguished benchmark weighting from underlying instability. For A2, the frozen-original, core-only, and regime-balanced summaries were similar: mean Δ*C* was +0.014, +0.019, and +0.016, while negative-transfer rates were 0.215, 0.210, and 0.219. For A3, stress-heavy weighting materially increased the magnitude of failure: mean Δ*C* changed from −0.025 in the frozen benchmark to +0.003 in the 36 core scenarios and −0.001 under regime-balanced weighting. Nevertheless, A3 negative-transfer rates remained 0.325 and 0.344 under the latter two summaries, with overall catastrophic-transfer frequencies of 0.153 and 0.174. Thus, the original stress composition amplified A3’s average and catastrophic failure but did not create the underlying negative-transfer instability (Supplementary Table S10).

The ordering was also stable to neighboring definitions of harm. Across Δ*C* ≤ −0.01, −0.02, −0.03, negative-transfer rates were 0.290/0.215/0.153 for A2 and 0.547/0.486/0.429 for A3. Across catastrophic definitions Δ*C* ≤ −0.04, −0.05, −0.06, the corresponding rates were 0.107/0.072/0.047 for A2 and 0.376/0.327/0.282 for A3. These analyses describe sensitivity of the estimated safety profile; they do not replace the frozen −0.02*/* −0.05 definitions or reopen the HOLD (Supplementary Table S11).

### 3.2. Training-set prognostic compatibility was identifiable but did not order transfer utility

The post-HOLD diagnostics distinguished training-set prognostic compatibility from the operational question of whether a transfer architecture would improve on a target-only model. At 29 target events, the frozen source risk evaluated against target-training outcomes separated the predefined mechanistically compatible regimes from incompatible regimes with AUROC 0.980. The same statistic performed below chance for classifying independent-test benefit versus harm: AUROC was 0.254 for A1 and 0.402 for A2. The prospectively frozen opening rule for the proposed A6 trust/abstention branch was therefore not met.

Numerically reversing the marginal A1 direction would yield AUROC 0.746 at 29 events, but this was observed post hoc and was not an eligible trust rule. More importantly, the A1 prognostic-compatibility–utility Spearman association changed from −0.385 marginally to +0.226 after removal of regime means; for A2 it changed from −0.168 to +0.027. Thus, the sub-chance marginal AUROC did not identify a stable inverse decision rule. Within the 22 mechanistically compatible scenarios, the median within-scenario association was positive (+0.355 for A1 and +0.250 for A2), with a positive sign in 21 of 22 scenarios for each model.

A comparator-free sensitivity reached the same qualitative conclusion. When high and poor absolute test discrimination were defined as *C* ≥ 0.55 and *C* ≤ 0.50, respectively, prognostic-compatibility AUROC was 0.203 for A1 and 0.262 for A2. Comparator decomposition also showed that the prognostic-compatibility statistic was negatively associated with both transfer-model discrimination (*ρ* = −0.604 for A1, −0.542 for A2) and B0 discrimination (*ρ* = −0.568). Together, these analyses show that the training-set statistic tracked generator structure but did not provide a single monotonic ordering of architecture-level held-out utility.

Useful transfer nevertheless existed within the evaluated model family. An outcome-aware oracle restricted to A1 achieved an equal-scenario mean gain of +0.036 over B0 with no negative transfer, and an oracle choosing per replicate among B0, A1, and A2 achieved +0.040, also with no negative transfer. These are optimistic upper bounds because independent-test outcomes were used for selection and evaluation. Among implementable event-count-only policies, A2 first met the 0.10 negative-transfer-rate criterion when transfer was restricted to settings with at least 20 target events; it was active in 50% of scenarios, with mean gain +0.008 and negative-transfer rate 0.092. This diagnostic did not establish satisfaction of the complete original protection rule.

### 3.3. Transfer utility was lowest under partial transportability

The controlled mechanism experiment evaluated 6,000 replicates generated from seeds disjoint from the 21,600 replicates used for architecture development. The experiment crossed four transport regimes with target event budgets of 10, 29, and 40. Head retargeting (M2) improved on the target-only model in the fully transportable, nontransportable, and misleadingsource regimes, whereas its mean gain was negative in the partially transportable regime at all three event budgets (Table 3; Fig. 3C).

**Table 3.** Mean Uno-C gain of head retargeting over the target-only model (Δ*C* = *C*_M2_ −*C*_M0_) in the controlled new-seed experiment. Each cell contains 500 replicates.

| Events | R0 full | R2 partial | R4 none | R5 reversal |
| --- | --- | --- | --- | --- |
| 10 | +0.0503 | −0.0068 | +0.0250 | +0.0717 |
| 29 | +0.0491 | −0.0083 | +0.0255 | +0.0588 |
| 40 | +0.0443 | −0.0140 | +0.0193 | +0.0497 |

**Figure 3.**
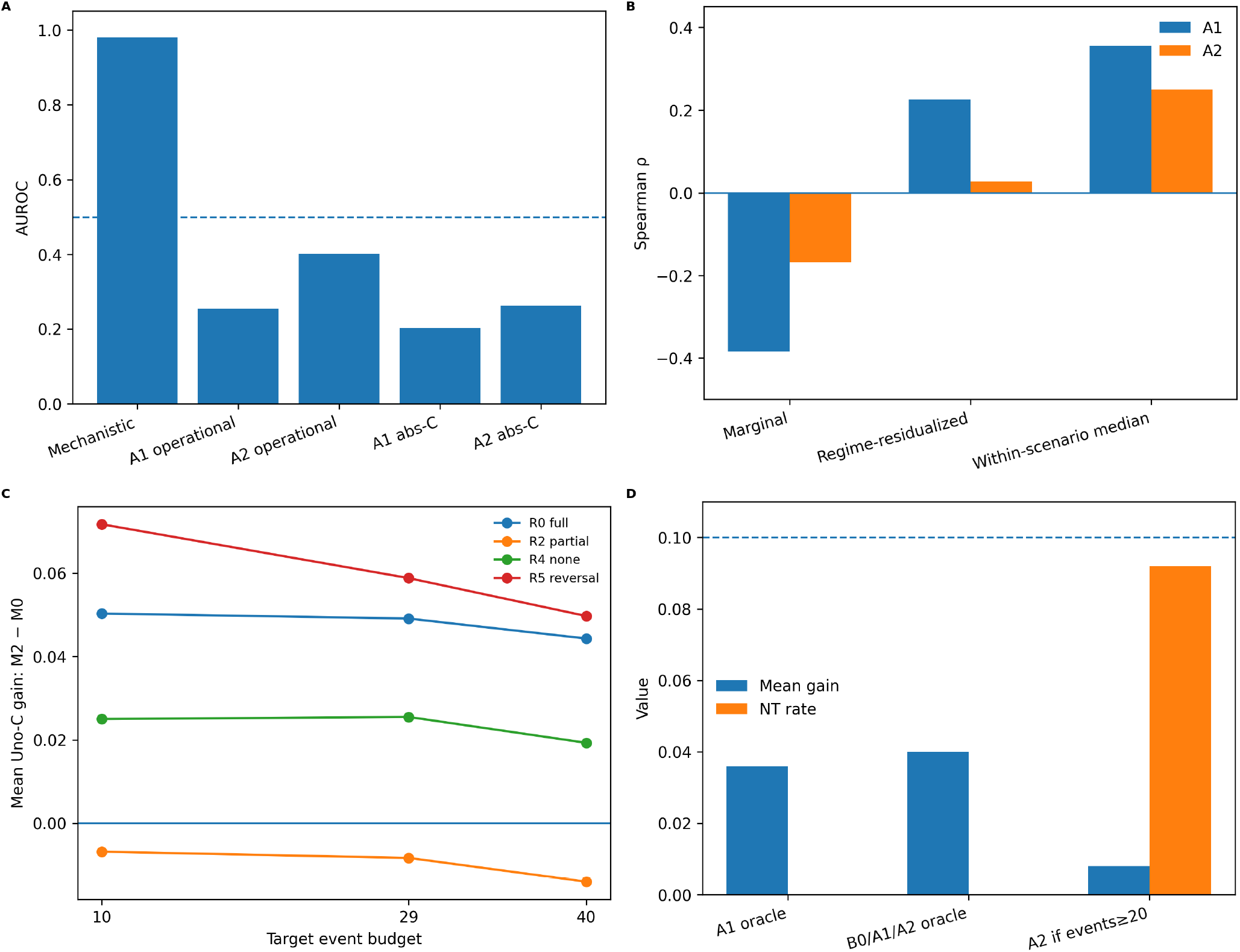
Training-set prognostic compatibility, transfer utility, and discrete regime structure. **A**, The same training-set prognostic-compatibility statistic strongly identified mechanistic generator regimes but performed below chance for the marginal operational benefit-versus-harm task. **B**, Marginal, regime-residualized, and within-scenario associations between training-set prognostic compatibility and transfer gain for A1 and A2. **C**, Mean Uno-C gain of head retargeting over the target-only model across the four predefined generator regimes and three event budgets; regimes are categorical and the connecting lines are used only to aid visual comparison. **D**, Outcome-aware oracle upper bounds and the fixed event-count abstention policy.

At the primary 29-event budget, all three contrasts against the partially transportable regime were positive, with bootstrap intervals excluding zero. The difference in transfer gain was +0.057 for R0 versus R2 (95% CI 0.051–0.064), +0.034 for R4 versus R2 (95% CI 0.028–0.040), and +0.067 for R5 versus R2 (95% CI 0.061–0.073). The frozen post-H1–H4 audit criterion for a discrete heterogeneity valley was therefore met.

The comparison with R4 constrains the interpretation of this pattern. R4 contained no transportable source effect, yet head retargeting performed better relative to B0 in R4 than in R2. A simple explanation in which transfer performance declines only as the amount of transportable source signal becomes smaller is therefore insufficient.

The four generator regimes jointly changed transportable fraction, effect concordance, and the fraction of sign-reversed nontransportable effects. The result consequently identifies a reproducible difficulty in the predefined partially transportable configuration rather than the isolated causal effect of one continuous heterogeneity parameter. The interpretation rule for this regime-shape audit was frozen after H1–H4 were known but before the corresponding regime-shape quantities were read (Table 1).

### 3.4. Target-head adaptation recovered global sign reversal through orientation alone

Seven model interventions were evaluated on the new simulation replicates, with four mechanism contrasts (H1–H4) specified before model fitting.

In the 29-event misleading-source cell, applying the frozen source network without target fitting produced a mean Uno-C of 0.340. Its negative-transfer and catastrophic-transfer rates were both 1.000. Re-estimating only the prognostic head while keeping the source encoder frozen increased Uno-C to 0.661; negative transfer fell to 0.022 and catastrophic transfer to 0.002. The paired Uno-C improvement over the unadapted source model was approximately +0.321, and the direction of the contrast was also positive at 10 and 40 events. H1 therefore satisfied its frozen support rule.

The later bounded sign-inversion audit showed that this recovery was almost entirely explained by orientation. At 29 events in R5, reversing the frozen source score changed its mean Uno-C from 0.340 to 0.660. The retargeted head achieved 0.661, only +0.0016 above the sign-reversed score (95% CI 0.0012–0.0019). The complete interval lay inside the frozen ±0.02 equivalence margin, giving the audit classification SIGN_INVERSION_EQUIVALENT. Patient-level risks from the retargeted and original source heads had a median correlation of −0.999.

The directional control behaved differently in the fully transportable regime. At 29 events, the median correlation between M2 and M1 risk scores was +0.998, and the adapted head retained positive alignment with the source head. Mean discrimination of the adapted head was slightly lower than that of the unadapted source model in this regime (0.639 versus 0.650), indicating that target adaptation was not uniformly beneficial even when source and target effects were highly concordant. No paired interval was computed for this contrast.

Head geometry changed more than prediction ranking in R5. The median cosine between the retargeted and source heads was −0.589, with a median orthogonal fraction of 0.808. These geometric quantities were secondary diagnostics and were not used to upgrade the prediction-level sign-equivalence classification.

Removing the penalty that centered the adapted target head on the source head did not produce a material discrimination gain. At 29 events in R5, the paired Uno-C difference between M2 and M3 was approximately −4 × 10^−6^ (95% CI approximately −1.7 × 10^−5^ to +4 × 10^−6^), far below the frozen +0.01 H2 requirement. Both secondary directions were non-worse: the paired IBS difference was approximately −3.7 × 10^−5^ and the paired held-out risk-score SD difference was 0.0022. H2 was therefore not supported because its primary discrimination criterion failed, despite both secondary criteria being satisfied.

Across the four 29-event regimes, mean discrimination of the unadapted source model ranged from 0.340 to 0.650, whereas the retargeted head ranged from 0.512 to 0.661. This compression of discrimination across regimes is descriptive; the prespecified H1 contrast concerned recovery specifically in R5.

### 3.5. Prediction-time hardening improved discrimination and reduced negative transfer while increasing risk-score dispersion

The learned soft-gating model M4 remained vulnerable in the primary 29-event misleading-source cell. Its mean Uno-C was 0.579, corresponding to a mean gain of −0.023 relative to the target-only model. Negative transfer occurred in 52.4% of replicates and catastrophic transfer in 8.2%.

M5 changed no fitted model parameter. It replaced the learned continuous gate by its 0.5-thresholded binary value only at prediction time and retained the M4 residual coefficients. Mean Uno-C increased to 0.625, negative transfer fell to 2.6%, and catastrophic transfer to 0.2%. The paired Uno-C gain over M4 was approximately +0.045, exceeding the frozen +0.02 primary H3 requirement.

The frozen H3 machine status nevertheless remained DOES_NOT_SUPPORT. The composite rule also required an absolute reduction of at least 0.10 in catastrophic-transfer rate. After H1–H4 were known, an arithmetic feasibility audit showed that the realized M4 control rate was 0.082, making a 0.10 absolute reduction impossible even if the treatment rate were zero. This did not alter the frozen classification: the primary H3 component was supportive, while the secondary component was structurally unassessable.

The truth-gated M6 intervention did not satisfy H4 on its primary criterion. Using the known transportability mask as a fixed binary gate and refitting the complementary residual gave a mean Uno-C of 0.591. The paired gain over M4 was approximately +0.011, below the frozen +0.02 requirement. H4 therefore remained unsupported independently of the same infeasible catastrophic-rate component.

**Figure 4.**
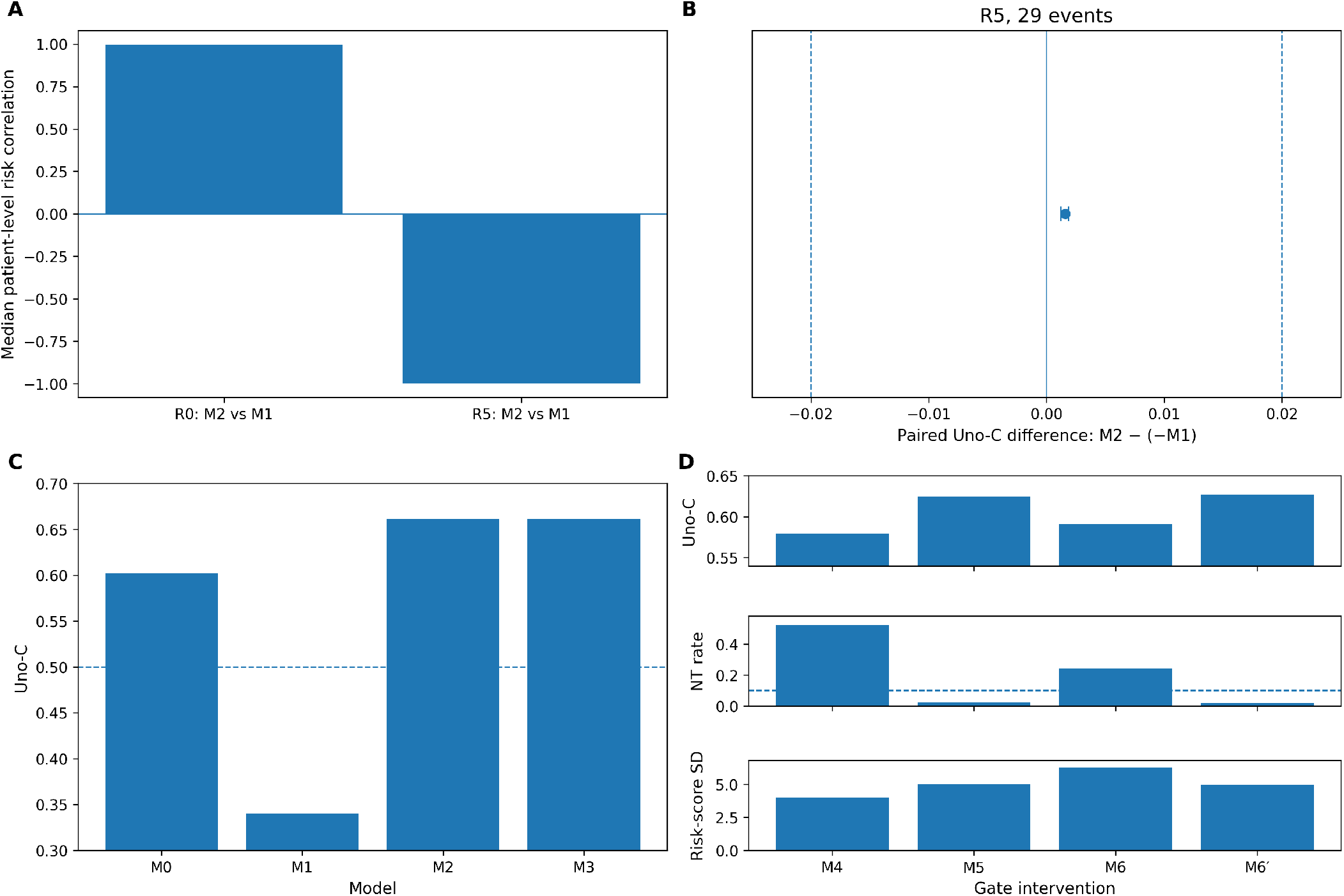
Controlled mechanism analyses on simulation seeds disjoint from those used for architecture development. **A**, Patient-level correlation between the frozen source risk and the retargeted-head risk in the fully transportable and misleading-source regimes. **B**, Paired difference between M2 and the sign-reversed frozen source score, −*M*1, in the primary R5 29-event cell, shown relative to the frozen ±0.02 sign-equivalence bounds. **C**, Discrimination of the target-only, frozen-source, retargeted-head, and no-centering models in the primary R5 29-event cell. **D**, Discrimination, negative-transfer rate, and risk-score dispersion for the learned soft gate M4, prediction-time hardened gate M5, oracle gate M6, and fixed-residual truth-gate diagnostic M6^′^.

A later diagnostic isolated gate substitution from residual refitting. M6^′^ replaced the M4 learned gate by the true binary transportability mask at prediction time while leaving the M4 residual unchanged. Mean Uno-C was 0.627, negative transfer 0.020, and catastrophic transfer 0.000. Relative to M4, its paired Uno-C gain was +0.0478 (95% CI 0.0457–0.0499). It exceeded M5 by only +0.0025 (95% CI 0.0017–0.0034).

This comparison is asymmetric: the M6^′^ residual had been optimized under the learned M4 soft gate rather than under the substituted true mask. M6^′^ is therefore a fixed-residual diagnostic and does not establish optimality of the learned versus true gate. It also cannot change the frozen H4 status.

The interventions also changed prediction error and riskscore scale. In the same cell, head retargeting M2 had IBS 0.199 and risk-score SD 0.83, compared with 0.315 and 4.12 for the target-only model and 0.328 and 4.02 for soft gating. Hardening the learned gate reduced IBS from 0.328 to 0.297 but increased risk-score SD from 4.02 to 5.02. Thus, prediction-stage hardening improved discrimination and the empirical negative-transfer rates without compressing the scale of the resulting risk score.

### 3.6. Cross-species molecular structure was heterogeneous across biological programs

The secondary outcome-blind biological analysis characterized structural preservation between DOG^2^ and TARGET expression without using survival outcomes or completed TARGET model results. Across the 50 frozen Hallmark gene sets, cross-cohort concordance of within-set gene–gene correlation structure was positive but heterogeneous. The median Spearman edge concordance was 0.319 (interquartile range 0.281– 0.399), with individual Hallmarks ranging from 0.159 to 0.605. Comparison with 1,000 exact-size expression- and variance-matched random panels per Hallmark provided a reference for the relative preservation of this structure. The median matched-control percentile across the Hallmark landscape was 0.783. Taken descriptively, the frozen Hallmark representation was therefore shifted toward the upper part of its matched-random reference distributions rather than being centered near the 0.50 percentile. This tendency was far from uniform: individual Hallmarks spanned almost the complete possible percentile range, from 0.005 to 1.000. Because Hallmark gene sets overlap, this landscape summary is descriptive and is not an inferential test across 50 independent modules (Fig. 5).

**Table 4.** Controlled model comparison in the primary 29-event misleading-source cell. Each row summarizes 500 new replicates. Δ*C* is relative to the target-only model M0. M6^′^ is a later bounded diagnostic and is not part of the frozen H1–H4 model set.

| Model | Definition | Uno-C | $\Delta C$ | IBS | Risk SD | NT rate | Catastrophic |
| --- | --- | --- | --- | --- | --- | --- | --- |
| M0 | Target-only | 0.602 | — | 0.315 | 4.12 | 0.000 | 0.000 |
| M1 | Frozen source network, zero-shot | 0.340 | −0.262 | 0.433 | 6.74 | 1.000 | 1.000 |
| M2 | Frozen encoder, retargeted head | 0.661 | +0.059 | 0.199 | 0.83 | 0.022 | 0.002 |
| M3 | M2 without source-head centering | 0.661 | +0.059 | 0.199 | 0.84 | 0.024 | 0.002 |
| M4 | Learned soft module gate | 0.579 | −0.023 | 0.328 | 4.02 | 0.524 | 0.082 |
| M5 | M4 gate hardened at prediction | 0.625 | +0.023 | 0.297 | 5.02 | 0.026 | 0.002 |
| M6 | True binary gate with refitted residual | 0.591 | −0.011 | 0.342 | 6.23 | 0.240 | 0.002 |
| M6' | True gate substituted into fixed M4 residual | 0.627 | +0.025 | 0.295 | 4.95 | 0.020 | 0.000 |

**Table 5.** Frozen artifact terminology and its scientific interpretation in the final analysis.

| Record | Frozen artifact label or rule | Scientific reading |
| --- | --- | --- |
| +0.02 | Positive operational reference used in post-HOLD diagnostics | A post-HOLD utility reference, not an original positive-utility threshold in the frozen A2/A3 architecture-selection rule. |
| beneficial_regime | Legacy artifact label applied to R0–R3 | R0–R3 are interpreted as mechanistically compatible generator regimes; the label does not assert empirical benefit of a particular transfer architecture. |
| H3 | DOES_NOT_SUPPORT | The primary Uno-C requirement was met, but the prespecified absolute catastrophic-rate reduction of 0.10 was structurally unattainable because the realized control rate was 0.082. The frozen machine status is retained. |
| H4 | DOES_NOT_SUPPORT | The primary Uno-C gain was approximately +0.011, below the frozen +0.02 requirement. The catastrophic-rate component was also structurally unassessable, but the primary criterion independently supports the negative H4 classification. |

**Figure 5.**
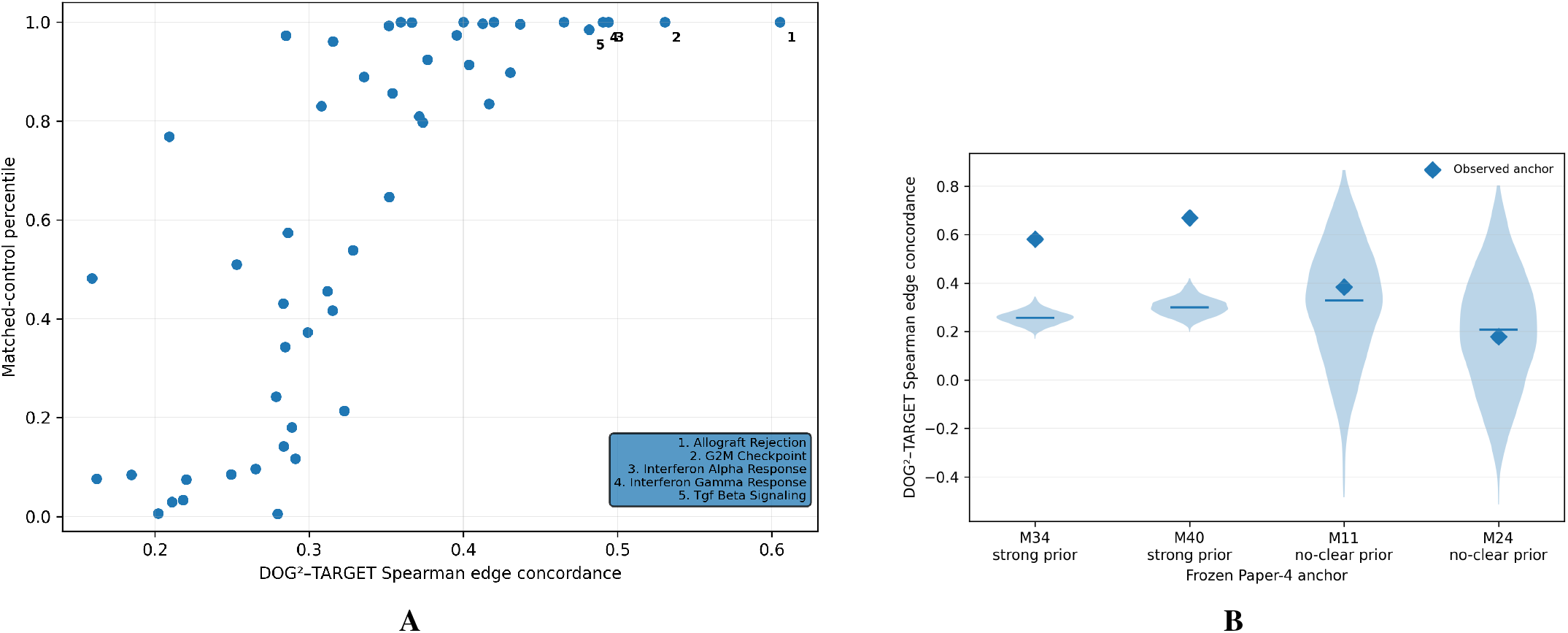
Outcome-blind cross-species structural preservation across the frozen biological representation. **A**, Structural concordance across all 50 Hallmark gene sets, shown against the percentile of 1,000 exact-size expression- and variance-matched random controls. Numbered points denote the five Hallmarks with the highest observed DOG^2^–TARGET Spearman edge concordance. **B**, Observed concordance of four independently frozen molecular-program anchors relative to their matched-control distributions. M34 and M40 had previously been classified as showing strong external canine representation preservation, whereas M11 and M24 had no clear external preservation.

The highest observed structural concordance occurred for allograft rejection (*ρ* = 0.605, matched-control percentile 1.000), G2M checkpoint (*ρ* = 0.531, percentile 1.000), interferon-alpha response (*ρ* = 0.494, percentile 1.000), interferon-gamma response (*ρ* = 0.490, percentile 1.000), and TGF-beta signaling (*ρ* = 0.482, percentile 0.985). These quantities describe preservation of within-program correlation structure and do not imply equivalent pathway activity or prognostic effects across species.

The four independently frozen molecular-program anchors provided a separate coherence check. These programs had been defined previously in an outcome-blind cross-cohort preservation analysis conducted independently of the present TARGET evaluation. M34 and M40 entered the current analysis with frozen labels indicating strong external canine representation preservation, whereas M11 and M24 entered with frozen labels indicating no clear external preservation. M34 and M40 had cross-species edge concordances of 0.581 and 0.670, respectively, and both occupied the 1.000 matched-control percentile. In contrast, M11 had *ρ* = 0.383 and a matched-control percentile of 0.594, whereas M24 had *ρ* = 0.178 and a percentile of 0.454. The anchors were retained regardless of whether the current structural statistic agreed with their prior labels.

The anchor analysis was descriptive and was not used to select biological programs or alter the transfer models. Likewise, the Hallmark-level landscape was not treated as 50 independent observations because the gene sets overlap. The result therefore provides biological context for a heterogeneous cross-species representation rather than an independent test that structural concordance predicts transfer utility.

### 3.7. Human TARGET evaluation remained unresolved under the frozen uncertainty rule

The frozen human evaluation included 86 TARGET osteosarcoma cases with 29 observed overall-survival events and 57 censored observations. The single censored zero-time case was retained unchanged. All nine frozen models produced valid Uno-C and IBS estimates in all 20 repeated cross-validation runs, and the patient-clustered bootstrap obtained 5,000 valid draws after 6,418 attempts. No model selection was performed. Discrimination and prediction-error estimates varied substantially across the frozen model registry (Table 6). The largest

**Table 6.** Frozen descriptive evaluation in 86 TARGET osteosarcoma cases (29 OS events). Uno-C intervals are from the prespecified patient-clustered bootstrap. IBS and risk-scale quantities are point estimates. All models had 20 of 20 valid repeated-CV estimates.

| Model | Definition | Uno-C [95% CI] | IBS | Risk SD | q99–q01 |
| --- | --- | --- | --- | --- | --- |
| T0 | Target-only ridge Cox | 0.5569 [0.4793, 0.6394] | 0.2093 | 3.799 | 13.960 |
| T1 | Frozen canine ridge Cox | 0.5956 [0.4998, 0.7078] | 0.1590 | 0.142 | 0.542 |
| T2 | Canine Cox + target residual | 0.5602 [0.4836, 0.6403] | 0.2038 | 4.599 | 16.570 |
| N0 | Frozen canine neural model | 0.5482 [0.4491, 0.6500] | 0.2366 | 7.084 | 23.059 |
| N1 | Frozen encoder + retargeted head | 0.5362 [0.4382, 0.6256] | 0.1691 | 0.384 | 1.410 |
| N2 | N1 without source centering | 0.5356 [0.4381, 0.6252] | 0.1691 | 0.384 | 1.410 |
| N3 | Rank-4 residual adapter | 0.5644 [0.4792, 0.6467] | 0.2046 | 1.214 | 4.441 |
| N4 | Target neural model from scratch | 0.6141 [0.5312, 0.6935] | 0.2388 | 7.397 | 23.484 |
| N5 | Full neural fine-tuning | 0.6199 [0.5272, 0.7091] | 0.2341 | 5.720 | 19.575 |
| *Frozen paired Uno-C contrasts* |  |  |  |  |  |
| N1–N0 |  | –0.0121 [–0.1301, +0.0998] |  |  |  |
| N0–T0 |  | –0.0086 [–0.1081, +0.0848] |  |  |  |
| N1–T0 |  | –0.0207 [–0.1307, +0.0770] |  |  |  |
| T1–T0 |  | +0.0387 [–0.0471, +0.1393] |  |  |  |
| N1–N2 |  | +0.0005 [–0.0005, +0.0021] |  |  |  |
| N3–T0 |  | +0.0075 [–0.0892, +0.0926] |  |  |  |

Uno-C point estimates were observed for the target-trained neural model N4 (*C* = 0.614, 95% CI 0.531–0.693) and the fully fine-tuned neural model N5 (*C* = 0.620, 95% CI 0.527– 0.709). No frozen paired contrast directly compared N4 with N5, so these point estimates do not establish a benefit of full fine-tuning over target-only neural training. Moreover, their prediction-error profiles were not favorable: N4 and N5 had IBS 0.239 and 0.234, respectively, versus 0.166 for the fold-local no-covariate Kaplan–Meier reference, with risk-score SDs of 7.40 and 5.72. Their higher discrimination point estimates therefore did not translate into uniformly favorable prediction-error or risk-scale behavior.

The frozen canine classical model T1 achieved an Uno-C of 0.596 (95% CI 0.500–0.708), compared with 0.557 (0.479– 0.639) for the target-only classical model T0. The paired T1–T0 difference was +0.039, but its 95% CI was wide and included zero (−0.047 to +0.139). T1 also had the lowest IBS among the nine models (0.159) and a markedly compressed held-out riskscore scale (SD 0.142; 99th–1st percentile range 0.542). These quantities are descriptive and were not used to select a model.

The post-opening no-covariate Kaplan–Meier reference contextualized the absolute IBS scale. Using the exact 20-by-5 frozen outer partitions and the same IBS time-grid, censoring, and aggregation mechanics, the fold-local training-only KM predictor had IBS 0.1656. T1 was the only frozen model with a lower IBS, and the difference was small (0.1590 versus 0.1656; ΔIBS = −0.0066). N1 and N2 were slightly above the null (both approximately 0.169), and the remaining models were farther above it. Because the KM reference assigns the same survival curve to every held-out patient within a fold, this comparison shows that a low absolute IBS can arise largely from cohort-level survival prediction and should not by itself be interpreted as strong individualized discrimination (Supplementary Table S12).

The outcome-zero-shot neural source model N0 had an Uno-C of 0.548 (95% CI 0.449–0.650). Retargeting only its prognostic head did not improve the discrimination point estimate: N1 had Uno-C 0.536, giving *N*1 −*N*0 = −0.012 (95% CI −0.130 to +0.100). Removing the source-head centering penalty had essentially no effect (*N*1 −*N*2 = +0.0005, 95% CI −0.0005 to +0.0021). The rank-4 residual adapter N3 had an Uno-C of 0.564, with *N*3 − *T*0 = +0.007 (95% CI −0.089 to +0.093).

The frozen point-estimate rules initially mapped the TARGET result to T-A: N0 was above 0.50, the N1–N0 difference was below +0.02, and the N0–T0 difference was greater than −0.02. The uncertainty rules, however, required this interpretation to be downgraded to T-D. The 95% bootstrap interval for N0 included 0.50 (0.449–0.650), leaving source-score orientation unresolved. In addition, both T-A-defining paired intervals spanned the complete frozen neutral region from −0.02 to +0.02: N1–N0 was −0.012 (95% CI −0.130 to +0.100), and N0–T0 was −0.009 (95% CI −0.108 to +0.085). The final frozen human interpretation was therefore T-D, indicating an unresolved descriptive pattern.

This result did not alter the simulation decision HOLD_NO_SAFE_SELECTABLE_ARCHITECTURE, reopen the A6 branch, or assign a synthetic generator regime to the human cohort. GSE21257 and GSE39055 outcomes remained sealed. TARGET therefore served only as the prespecified descriptive human stress test of the frozen model registry.

## 4. Discussion

This study examined cross-species survival transfer under severe target-event scarcity using a chronologically separated evaluation design. Neither architecture eligible for selection satisfied the frozen negative-transfer protection rule. This negative result was interpretable because post-HOLD oracle analyses showed that the 0.10 negative-transfer threshold was attainable within the same known-truth benchmark. A3 also satisfied its prespecified module-recovery criterion while still failing the protection rule. Recovering which simulated modules were transportable was therefore not sufficient to guarantee safe borrowing by the evaluated architecture.

The post-HOLD robustness summaries sharpen this conclusion. Across the complete frozen model family, the architecture with the highest mean Uno-C was not the one with the lowest negative-transfer risk, illustrating why average performance alone was an inadequate selection target. Alternative benchmark weighting had little effect on A2 but substantially attenuated the magnitude of A3’s average and catastrophic failure. Even after restricting to the 36 core scenarios or balancing all six regimes equally, however, A3 negative-transfer rates remained above 0.32. Neighboring harm definitions also preserved the same qualitative ordering. The frozen stress-heavy benchmark therefore magnified A3’s failure but did not manufacture it.

The results also separate training-set prognostic compatibility from transfer utility. The prognostic-compatibility statistic identified the predefined generator regimes with high accuracy but performed below chance for the marginal operational benefit-versus-harm task. Although post-hoc reversal of the A1 direction would numerically give AUROC 0.746 at 29 events, the association itself changed sign after conditioning on regime: the marginal A1 association of −0.385 became +0.226 after regime residualization. The statistic therefore contained information about the source–target setting without providing a reliable monotonic trust rule for the evaluated transfer procedures.

Consistent with that distinction, the prospectively gated A6 feasibility branch did not meet its frozen opening criterion.

The controlled experiment on disjoint simulation seeds localized two distinct mechanisms. In the misleading-source regime, retargeting only the prognostic head produced a large improvement over the frozen source predictor, but the later sign-inversion audit showed that almost all of the recovered ranking was already obtainable by reversing the orientation of the source score. This was therefore a comparatively simple orientation correction rather than evidence that the adapted model recovered substantial new ranking structure. In contrast, transfer utility was lowest in the partially transportable regime and remained negative at all three evaluated event budgets. The nontransportable regime performed better than this partial regime, ruling out a simple monotonic explanation based only on the amount of transferable source signal. Because the predefined regimes changed several generator properties jointly, the observed valley should be interpreted as a reproducible difficulty of the partial-transport configuration rather than the isolated causal effect of one continuous heterogeneity parameter.

Prediction-time gate hardening provided the clearest implementable positive intervention. Replacing the learned continuous borrowing gate by a fixed binary threshold at prediction time required no refitting, improved discrimination, and markedly reduced the observed rates of negative and catastrophic transfer in the misleading-source cell. Its prespecified composite safety criterion was nevertheless not met because the required absolute catastrophic-rate reduction was mathematically unattainable at the realized control rate. The frozen classification was therefore retained. Hardening also increased riskscore dispersion even while improving discrimination and IBS. This divergence shows why transfer behavior cannot be summarized by concordance alone: ranking, prediction error, and score scale can move in different directions.

The secondary outcome-blind biological analysis placed the computational findings in the real canine–human expression setting rather than serving as an independent validation of transfer performance. Previous studies have reported conserved transcriptional and microenvironmental structure between canine and human osteosarcoma [2, 3, 5, 6]. Across the full frozen Hallmark representation, structural preservation was heterogeneous rather than uniformly high or low, and matched-control percentiles extended across nearly their full possible range. Four pre-existing coherence anchors from an earlier outcome-blind analysis by the same authors were retained without reselection: M34 and M40, previously labeled as strongly preserved, occupied the extreme upper tail of their matched-control distributions, whereas M11 and M24 did not. Because the structural statistic measures expression organization of the type that the transfer representation itself can exploit, this analysis characterizes the biological domain-shift setting and does not establish that structural concordance independently predicts prognostic transfer utility.

TARGET-OS was assigned a deliberately different evidentiary role. With only 29 observed OS events, the cohort was not used to select an architecture or to retrospectively confirm the simulation mechanisms. The nine frozen comparators were instead evaluated as a descriptive human stress test under fixed preprocessing, resampling, and reporting rules.

In TARGET-OS, the frozen model registry produced heterogeneous descriptive results rather than a single transferable pattern. N4 and N5 had the largest Uno-C point estimates, but both had substantially higher IBS than the fold-local no-covariate Kaplan–Meier reference and large risk-score dispersion; their discrimination advantage therefore did not translate into uniformly favorable prediction-error behavior. Conversely, the frozen canine classical model T1 had the lowest IBS, but its improvement over the no-covariate reference was only 0.0066 (0.1590 versus 0.1656). These contrasts prevent either a high Uno-C point estimate or a low absolute IBS from being interpreted in isolation as strong individualized prognostic performance. Uncertainty in Uno-C remained substantial, and the frozen T-A–T-D rules ultimately assigned T-D. The human analysis therefore remained descriptively unresolved and did not alter the simulation HOLD.

Whatever ordering is observed in TARGET cannot change the frozen simulation decision. This separation is particularly important in rare-disease settings, where a single small cohort can otherwise become both the development set and the evidence used to justify development choices.

Several limitations define the scope of these conclusions. First, all primary mechanism results arise from one known-truth generator family. The benchmark was deliberately stress weighted: R2 and R5 each contributed 78 of the 180 scenarios because nuisance-axis stress tests were concentrated in these two regimes, whereas each remaining regime contributed six core scenarios. The post-HOLD core-only and regime-balanced sensitivities showed that this composition amplified the magnitude of A3 failure, especially its mean Δ*C* and catastrophic-transfer frequency, while leaving A2 comparatively stable. A3 nevertheless retained negative-transfer rates above 0.32 under both alternative estimands. The frozen equal-scenario aggregate should therefore be read as behavior under a stress-heavy benchmark rather than an expected negative-transfer probability in an unselected application, while the sensitivity results indicate that the underlying instability was not solely a weighting artifact. The controlled regimes also changed multiple transport properties jointly, which prevents causal attribution of the partial-transfer valley to one parameter.

Second, the real-data evaluation remains small and retrospective. Repeated cross-validation and patient-clustered bootstrap uncertainty in TARGET do not replace external prospective validation. The biological analysis has a weaker evidentiary status because it was frozen after TARGET outcome access, although before the completed TARGET results were inspected, and overlapping Hallmark sets prevent treating its 50 modules as independent observations.

Third, calibration slope and calibration-in-the-large were evaluated as secondary simulation diagnostics, but neither entered the frozen architecture-selection rule. Meeting a discrimination-based transfer-safety criterion would therefore not by itself establish calibrated absolute risk or clinical utility; the full calibration results are reported in Supplementary Table S3. Similarly, the study evaluates protection against negative transfer, not treatment benefit or decision-analytic usefulness.

Finally, the frozen representation used independent within-domain standardization rather than a learned covariance-alignment method. More elaborate approaches such as CORAL [14] could in principle be estimated fold-locally without held-out leakage, but they were not part of the frozen model registry and adding them after the benchmark results were known would have introduced a new post-result degree of freedom. They would also require additional regularization in the present setting: a typical TARGET outer-training fold contains only about 69 patients for 50 Hallmark dimensions, making direct covariance estimation unstable and introducing a further shrinkage choice. Evaluation of such alignment methods therefore requires a separately prespecified study rather than posthoc extension of the current benchmark.

The broader implication is that cross-domain biomedical learning should separate three questions: whether outcome-blind source and target structure is concordant, whether training-set prognostic compatibility is informative, and whether a particular model benefits from borrowing and can do so safely using the target information available. These questions were not interchangeable in the present experiments. Known-truth simulation, explicit negative-transfer criteria, and chronological separation of model development from later diagnostics provide one way to evaluate them before a small human cohort is used for descriptive assessment.

## 5. Conclusion

Cross-species molecular transfer under severe target-event scarcity requires separating training-set prognostic compatibility, outcome-blind structural concordance, and the utility and safety of a specific transfer architecture. In a prespecified known-truth benchmark, neither selectable architecture satisfied the frozen negative-transfer protection rule, although oracle analyses showed that the protection threshold was attainable within the same benchmark. Controlled new-seed experiments further showed that global sign reversal could be corrected largely through score orientation, whereas partial transportability remained a distinct failure setting. Prediction-time hardening of a learned soft borrowing gate improved discrimination and substantially reduced empirical negative transfer without additional fitting, although its prespecified composite safety criterion was not met and risk-score dispersion increased. Post-HOLD weighting and definition sensitivities showed that the magnitude of failure depended partly on benchmark composition while the qualitative safety ordering persisted. In TARGET, the only IBS improvement over a fold-local no-covariate Kaplan–Meier reference was small, reinforcing the distinction between favorable absolute prediction-error values and individualized prognostic discrimination. These results support evaluating cross-domain transfer with explicit safety criteria and chronologically separated evidence before small human cohorts are used to judge model performance.

## Ethics Statement

This study performed secondary computational analyses of previously collected canine and human molecular datasets. No new human participants or animals were recruited, no intervention was performed, and no new biological specimens were collected as part of this study. Ethical approvals, consent procedures, and data-access conditions for the original cohorts are described in the corresponding source studies and data repositories.

## Code Availability

Code for cross-species feature construction, simulation generation, model fitting, frozen architecture evaluation, controlled mechanism experiments, biological-context analysis, and descriptive TARGET evaluation will be made publicly available with the final publication.

The release will include versioned analysis scripts, frozen configuration files, random-seed registries, model and metric definitions, analysis contracts, and machine-readable audit artifacts required to reproduce the reported results. Technical amendments are retained as separate versioned artifacts rather than replacing earlier records. A persistent public repository identifier will be added to the accepted manuscript.

## Data Availability

The canine DOG^2^ osteosarcoma transcriptomic dataset analyzed in this study is publicly available through the Gene Expression Omnibus under accession GSE238110. The human GSE16091 dataset is also available through the Gene Expression Omnibus. Human osteosarcoma molecular data used for the descriptive TARGET analysis were obtained from the NCI Therapeutically Applicable Research to Generate Effective Treatments (TARGET) osteosarcoma project.

All cross-species gene mappings, Hallmark gene-set memberships, analysis rosters, and derived non-identifying intermediate data required to reproduce the reported analyses will be released with the accompanying code repository, subject to the redistribution terms of the original data sources. Data from the original repositories will not be redistributed where the source terms require users to obtain them directly.

## Supplementary material

Supplementary Methods, Tables S1–S13, and Figures S1–S7 are provided as a separate supplementary file.

## Declaration of generative AI and AI-assisted technologies in the manuscript preparation process

During the preparation of this work, the authors used ChatGPT (OpenAI) and Claude (Anthropic) to assist with manuscript drafting, language refinement, and code-debugging discussion. After using these tools, the authors reviewed and edited the content as needed and take full responsibility for the content of the published article.

