## Supplementary material for "When Prognostic Compatibility Does Not Guarantee Transfer Utility in Event-Limited Cross-Species Survival Modeling": Suplimental File

---

**Supplementary Methods**

- Full chronological analysis and outcome-access firewall.
- Complete simulation generator and scenario construction.
- Frozen architecture-selection procedure and candidate registry.
- Full post-HOLD diagnostic sequence and stop rules.
- Controlled new-seed experiment and bootstrap procedures.
- TARGET endpoint reconciliation and post-opening technical amendment.
- Byte-level runner provenance and reproducibility checks.
- Outcome-blind biological-context matching procedure.
- Post-HOLD weighting, harm-definition, and TARGET KM-null sensitivity procedures.

**Supplementary Tables**

- Table S1. Frozen model and hyperparameter registry.
- Table S2. Complete simulation-generator specification.
- Table S3. Secondary simulation metrics, including calibration slope and calibration-in-the-large.

- Table S4. Full post-HOLD diagnostic sequence and evidence status.
- Table S5. Exact H1-H4 mechanism contrasts and frozen support rules.
- Table S6. Biological-context matching, Hallmark overlap, and matched-control results.
- Table S7. Complete TARGET fold-, repeat-, bootstrap-, and model-level results.
- Table S8. Software environment, analysis chronology, and major frozen-artifact SHA-256 ledger.
- Table S9. Descriptive all-model simulation safety summary.
- Table S10. Frozen-original, core-only, and regime-balanced aggregation sensitivity.
- Table S11. Neighboring negative- and catastrophic-transfer definition sensitivity.
- Table S12. Fold-local no-covariate TARGET Kaplan–Meier IBS reference.
- Table S13. Chronology and provenance of pre-existing biological anchors.

**Supplementary Figures**

- Figure S1. Complete simulation scenario composition.
- Figure S2. Event-budget-specific negative-transfer profiles.
- Figure S3. Comparator decomposition and within-scenario compatibility diagnostics.
- Figure S4. Event-count abstention ladder and oracle upper bounds.
- Figure S5. Complete H1–H4 secondary metrics and risk-scale diagnostics.
- Figure S6. Additional biological-context matched-control diagnostics.
- Figure S7. TARGET repeat-level distributions and sensitivity plots.

### Supplementary Methods

*S1. Analysis chronology and evidence governance*

The analysis was implemented as a sequence of versioned stages with explicit restrictions on what information could be read and what decisions could be changed at each stage. The purpose of this structure was to prevent reserved human outcomes or later simulation results from being used to modify earlier model definitions, thresholds, or interpretation rules.

The known-truth simulation design, candidate model registry, negative-transfer definitions, architecture-selection thresholds, and practical tie rule were frozen before any simulation model was fitted. GSE16091 had already been used as a sacrificial human premise cohort, but its numerical performance was explicitly prohibited from determining the simulation grid, architecture hyperparameters, or selection thresholds. TARGET-OS, GSE21257, and GSE39055 outcomes remained closed during this stage.

The initial simulation model matrix and the corresponding survival metrics were then evaluated under the frozen architecture-selection rule. Once this rule returned no selectable A2/A3 architecture, the decision was retained. Later analyses of the original 21,600 simulation replicates were classified as post-HOLD diagnostics. They could test implementation integrity, criterion attainability, compatibility signals, and explanatory mechanisms, but could not alter the original architecture decision or promote a new candidate into the selection set.

A separate controlled mechanism experiment was subsequently frozen before fitting any of its models. Its 12 cells, 6,000 new seeds, seven model interventions, and four H1–H4 contrasts were fixed prospectively. All 6,000 seeds were unique and had zero overlap with the seeds used in the original 21,600-replicate benchmark.

After H1–H4 had been evaluated, a final bounded simulation audit was specified to address sign inversion, discrete regime shape, and a fixed-residual truth-gate substitution. The rules for these new audit quantities were frozen before those quantities were read, but after H1–H4 were already known. Accordingly, this audit has a weaker evidentiary status than H1–H4 and cannot rewrite them. It also imposed a stop rule prohibiting additional simulation model branches or diagnostics from the same results before manuscript drafting.

The TARGET expression representation, clinical schema, repeated cross-validation mechanics, bootstrap procedure, model implementation sources,

DOG[2] source artifacts, output schema, and complete evaluation runner were then frozen before TARGET survival values were opened. TARGET was designated a descriptive, non-confirmatory analysis: it could not select an architecture, change the original HOLD, or reopen the A6 trust branch.

Finally, a secondary outcome-blind biological-context analysis was frozen after TARGET outcome access but while the TARGET model evaluation was still running and before its completed model results were inspected. This analysis therefore has the explicit status *post-TARGET-opening, pre-TARGET-result-inspection, outcome-blind*. It cannot be described as a pre-TARGET confirmatory analysis.

Each freeze generated machine-readable contracts and SHA-256 hashes. Downstream stages verified the required hashes before reading their scientific inputs. When a technical defect was identified after a freeze boundary, it was handled through a new versioned amendment rather than by silently changing the original artifact. Scientific status and technical execution status were recorded separately.

*S2. Known-truth simulation generator*

The simulation generator operated directly in a 50-dimensional module space. Ten of the 50 modules were causal in each replicate. Causal modules were sampled without replacement independently for each replicate.

The source population contained 186 observations and exactly 124 observed events. The source covariance matrix contained five blocks of 10 modules. Pairwise correlation was 0.30 within a block and 0.03 between modules belonging to different blocks. The matrix was projected to a valid positive-definite correlation matrix using a minimum eigenvalue of $10^{-5}$.

For each replicate, non-zero source Cox coefficients had independent magnitudes

$$|\beta_{S,j}| \sim U(0.20, 0.45) \tag{1}$$

and independent random signs. Synthetic event times followed a proportional-hazards model with baseline hazard 0.01. The linear predictor was clipped to $[-8, 8]$ for numerical stability.

Target covariance shift was generated independently of target outcome-effect transport. A random eight-dimensional low-rank covariance structure was converted to a correlation matrix and blended with the source correlation matrix,

Table 1: Frozen source–target transport regimes in the known-truth generator.

| ID | Regime | Transportable fraction | Effect-retention parameter | Nontransport |
|---|---|---|---|---|
| R0 | Fully transportable | 1.00 | 0.90 | [illegible] |
| R1 | Mostly transportable | 0.75 | 0.75 | [illegible] |
| R2 | Partially transportable | 0.50 | 0.50 | [illegible] |
| R3 | Weakly transportable | 0.25 | 0.25 | [illegible] |
| R4 | Nontransportable | 0.00 | 0.00 | [illegible] |
| R5 | Misleading source | 0.00 | −0.50 | [illegible] |

$$\Sigma_T = (1 - s)\Sigma_S + s\Sigma_{\text{random}}, \tag{2}$$

where the frozen shift strengths were $s \in \{0, 0.25, 0.50\}$.

The six predefined transfer regimes are shown in Table 1. A transportable mask was sampled over all 50 modules according to the frozen transferable fraction. For causal modules that were also transportable, the target coefficient retained the source direction and was multiplied by a normally distributed scale centered at the regime's effect-retention parameter with standard deviation 0.10 and lower bound 0.05. For causal modules classified as nontransportable, a prespecified fraction had its source direction reversed with approximately unit magnitude; remaining nontransportable causal effects were set to zero.

The module-level evolutionary prior was generated independently of survival outcomes after the transportability mask had been drawn. Under the correct prior, transportable modules were sampled from $\text{Beta}(8, 2)$ and nontransportable modules from $\text{Beta}(2, 8)$. The uninformative prior used $\text{Beta}(5, 5)$ for all modules. The misleading prior reversed the two correct-prior distributions.

Mapping corruption was outcome-independent. A frozen fraction of module correspondences was selected for corruption. Approximately half of the selected indices were dropped completely and represented by a mapping value of $-1$; the remaining corrupted indices were cyclically mis-mapped. Mapping-error fractions were 0, 0.10, and 0.30.

Target training sets were defined by an exact number of observed events rather than by a fixed number of patients. For a requested event count $E$ and target censoring fraction $c$, the target training size was

$$n_T = \max\left\{\text{round}\left(\frac{E}{1-c}\right), E+1\right\}. \tag{3}$$

Independent exponential censoring variables were rescaled so that the requested training event count was obtained exactly. No empirical survival times from any real cohort were used in this calibration. Each replicate additionally contained an independent target test population of 500 observations generated from the same target covariance and target coefficient vector. Target test event counts reflected the scenario-specific censoring fraction.

The primary event-count grid was $E \in \{5, 10, 15, 20, 29, 40\}$. The core phase diagram crossed all six event budgets with all six transport regimes, giving 36 core scenarios with 200 replicates each. Core scenarios used moderate covariance shift ($s = 0.25$), 40% censoring, no mapping error, and the correct source prior.

Stress scenarios were concentrated on R2 and R5 and on event budgets 10, 15, 29, and 40. For each event-budget/regime pair, one family crossed three covariance-shift levels with three censoring levels, and a second family crossed three mapping-error levels with three source-prior states. This produced 144 stress scenarios with 100 replicates each. The complete benchmark therefore contained

$$36 \times 200 + 144 \times 100 = 21{,}600 \tag{4}$$

replicates across 180 scenarios.

Because the stress grid intentionally concentrated on R2 and R5, these two regimes each contributed 78 of the 180 scenarios, whereas R0, R1, R3, and R4 each contributed six core scenarios. Scenario-level equal weighting was used for the frozen architecture-selection summaries.

Each replicate had a deterministic seed derived from the frozen base seed, scenario index, and replicate index. The simulation materialization stage saved the seeds, source and target coefficient truth, transportability masks, prior scores, and mapping-corruption maps rather than persisting all full matrices. The full source and target matrices were regenerated on demand by the SHA-locked authoritative generator. Six representative full-data fixtures were stored and required to reproduce exactly before the model-fitting stage could proceed.

Table 2: Frozen simulation model and optimization registry.

| Component | Main setting | Epochs | Learning rate | Regularization |
|---|---|---|---|---|
| Source linear Cox | $\alpha = 1.0$ | 120 | 0.05 | L2 |
| B0 target-only Cox | $\alpha = 2.0$ | 120 | 0.05 | L2 |
| B4 residual Cox | $\alpha = 2.0$ | 120 | 0.05 | L2 toward source |
| Source MLP | 50–32–16–1 | 120 | 0.010 | weight decay $10^{-4}$ |
| A0 | target neural scratch | 140 | 0.010 | weight decay $10^{-3}$ |
| A1 | frozen encoder + head | 80 | 0.020 | head-to-source L2 1[illegible] |
| A2 | latent residual adapter | 100 | 0.020 | adapter L2 $10^{-2}$ |
| A3 | module-selective borrowing | 120 | 0.020 | prior 0.10; gate L2 0[illegible] |
| A4 | full fine-tuning | 80 | 0.003 | weight decay $5 \times 10$[illegible] |

*S3. Frozen simulation-model implementation*

The simulation model matrix contained two classical anchors, five main neural models, and five A3 ablations. Model fitting used only synthetic data and did not read any DOG[2] or human expression or outcome values.

B0 was a target-only linear Cox model. B4 used the same target linear Cox objective with an L2 penalty centered on the mapped source coefficient vector. The neural source model contained 50 inputs, a 32-unit hidden layer, a 16-dimensional latent layer, hyperbolic-tangent activations, and a linear survival-risk head.

A0 trained this neural architecture from scratch on the target. A1 froze the source encoder and updated only a source-initialized prognostic head. A2 froze the source model and added a rank-4 residual adapter in latent space. A3 froze the source network and fitted one continuous borrowing gate per module together with a complementary linear target residual. A4 initialized the full network from the source model and updated all parameters.

The five A3 ablations removed the evolutionary prior, prevented target-driven gate adjustment, permuted the prior, permuted the source–target mapping, or permuted source survival outcomes. The model family was closed before the simulation matrix was fitted.

All simulation hyperparameters were fixed globally rather than tuned by scenario. Neural optimization used Adam [1], gradient-norm clipping at 5, fixed epoch counts, and no early stopping. The complete registry is shown in Supplementary Table S1.

Source pretraining was shared within each replicate across models that required the same source network. Source and target module matrices were standardized independently within their respective training domains. All fitted parameters and held-out risk predictions needed for later survival metrics were stored in restart-safe scenario shards.

*S4. Survival metrics and frozen architecture-selection rule*

The primary discrimination metric was Uno's IPCW concordance statistic [2], implemented with scikit-survival [3]. Larger risk scores represented higher hazard. For each synthetic replicate, the Uno-C evaluation horizon was the 80th percentile of target-training event times and was truncated strictly below the common observed train/test follow-up support. The same horizon was used for all models within a replicate.

Integrated Brier score was calculated using survival probabilities obtained from each model's saved target-training and target-test risk scores [4]. A Breslow baseline cumulative hazard [5] was estimated using target-training outcomes. The IBS grid contained 20 equally spaced time points from the 20th to 80th percentile of target-training event times after restriction to common train/test support.

Calibration slope was estimated by a one-dimensional Cox regression of the test outcome on the unchanged saved risk score. Calibration-in-the-large was computed at the median target-training event-time horizon as the difference between the complementary-log-log transformation of observed Kaplan–Meier test survival and that of mean predicted test survival. Ideal values were one for the slope and zero for the intercept. These quantities were descriptive and did not enter the A2/A3 architecture decision. Complete calibration summaries are reported in Supplementary Table S3.

For each replicate and model,

$$\Delta C = C_{\text{model}} - C_{\text{B0}}. \tag{5}$$

Negative transfer was defined as $\Delta C \leq -0.02$, and catastrophic negative transfer as $\Delta C \leq -0.05$.

For models with module-level borrowing gates, recovery of the known transportability mask was measured by AUROC of the continuous gate against the 50-dimensional transportability truth. False borrowing was the fraction of truly nontransportable modules with borrowing weight at least 0.5.

Metrics were first aggregated across replicates within a frozen scenario. Selection-level summaries then assigned equal weight to each of the 180 scenarios, rather than weighting scenarios by their different replicate counts.

Only A2 and A3 were eligible for final architecture selection. A candidate had to satisfy the aggregate negative-transfer rate limit of 0.10 and catastrophic negative-transfer rate limit of 0.05 separately in misleading-source and severe covariance/mapping-shift scenarios. A3 additionally required mean module-recovery AUROC of at least 0.70 across the partially transportable regimes and mean false borrowing no greater than 0.20 under R5.

If only one candidate satisfied all applicable protection criteria, that candidate would be selected. If neither did, the frozen result was `HOLD_NO_SAFE_SELECTABLE_ARCHITEC`[illegible] If both passed, mean Uno-C was primary. An absolute difference below 0.01 was treated as a practical tie, after which common-replicate IBS was compared. If the IBS difference was also below 0.01, the simpler A2 architecture was selected. Calibration was not a tie-breaking quantity.

*S5. Post-HOLD implementation and threshold diagnostics*

Once the original A2/A3 rule had been applied, subsequent analyses of the 21,600-replicate benchmark were explicitly post-HOLD. They could not alter the selection rule or make an additional architecture eligible.

The first bounded audit replayed structural invariants, saved model outputs, and the frozen selection summaries. A separate precision audit reconstructed Uno-C directly from the stored float32 predictions to verify that numerical serialization had not changed the frozen metric interpretation.

Threshold attainability was then examined with deliberately optimistic oracle policies. A true-target-risk oracle used known synthetic target risk to determine whether source transfer should be used. Regime-aware abstention policies used known generator regime membership. These policies were not considered implementable estimators; their sole purpose was to determine whether the frozen negative-transfer limit of 0.10 could in principle be achieved within the existing simulation environment.

A training-set prognostic-compatibility statistic was next defined. A frozen source linear Cox score was applied to target-training features after target-training-only standardization. No target outcome was used to refit the source score. Compatibility was the Uno-C of this fixed source score on the target-training patients themselves,

$$C_{\text{compat}} = C_{\text{Uno}}\left(r_{\text{source}}, Y_{\text{target,train}}\right). \tag{6}$$

Two distinct labels were evaluated. The secondary mechanistic label classified R0–R3 as compatible and R4–R5 as incompatible. The primary operational label was model-specific: an A1 or A2 replicate was labeled beneficial if independent test $\Delta C \geq +0.02$, harmful if $\Delta C \leq -0.02$, and neutral otherwise. Neutral replicates were excluded from operational AUROC.

Before computing these AUROCs, the A6 feasibility decision was frozen. At the primary 29-event budget, A6 could be opened for a new prospective contract only if operational AUROC was at least 0.70 for A1 or A2. A limited-future-method state required an AUROC between 0.65 and 0.70 at 29 events together with an AUROC of at least 0.70 at 40 events. All other outcomes closed this prospective branch for the present study. Even a GO state would only have authorized writing a new prospective contract; it would not have authorized immediate human outcome fitting.

A trivial event-count abstention family was also evaluated. For each $N \in \{5, 10, 15, 20, 29, 40, 41\}$, the policy used the frozen A1 or A2 predictor only when the target event count was at least $N$ and otherwise returned the exact B0 target-only model. $N = 41$ represented complete abstention. This diagnostic evaluated the aggregate negative-transfer-rate component and did not replace the full original protection rule.

Subsequent bounded audits examined the direction of the prognostic-compatibility signal, regime-residualized prognostic-compatibility–utility associations, within-scenario associations, decomposition of prognostic-compatibility relationships with absolute transfer-model and B0 concordance, and an absolute-discrimination sensitivity analysis. The absolute-C sensitivity labeled independent-test concordance $\geq 0.55$ as high and $\leq 0.50$ as poor. These analyses were post-HOLD and could not reopen A6.

Outcome-aware upper bounds were also computed by choosing between already evaluated predictors using the same synthetic test outcome on which the chosen prediction was evaluated. These quantities were explicitly treated as optimistically biased oracle bounds and not as implementable selection procedures.

*S6. Controlled new-seed mechanism experiment*

A separate mechanism experiment was frozen after the post-HOLD diagnostic sequence. It crossed event budgets 10, 29, and 40 with R0, R2, R4, and R5, giving 12 cells. Covariance shift was fixed at zero, censoring at 0.40, mapping error at zero, and the source prior at its correct state. Each cell contained 500 replicates and an independent target test population of 500 observations.

The 6,000 replicate seeds were materialized before model fitting. All seeds were unique and were checked against the complete original 05b seed universe; the overlap count was required to be zero. Each cell also underwent one deterministic smoke replay through the original authoritative 05b generator before model fitting.

Seven interventions were fitted by importing the original 05c implementation rather than recreating it:

- M0: exact target-only B0 replay;
- M1: frozen source neural encoder and source head, with no target fitting;
- M2: exact A1 source-centered target-head adaptation;
- M3: M2 with only the L2 penalty toward the source head set to zero;
- M4: exact A3 learned continuous gate and complementary residual;
- M5: the fitted M4 gate thresholded at 0.5 only at prediction time, with the M4 residual retained and no new fit;
- M6: the true synthetic transportability mask used as a fixed binary gate, with only the complementary residual fitted; this is a nonimplementable oracle.

All other source-model, mapping, optimization, and prediction functions were reused from the SHA-verified 05c implementation.

The four mechanism contrasts were frozen before any 05e2 prediction array was read. Their exact rules are given in Supplementary Table S5.

Uno-C used the same 80th-percentile training-event time-support rule as the original simulation benchmark. Secondary quantities were IBS, held-out population risk-score SD, the 99th-to-1st percentile risk range, negative-transfer rate, and catastrophic-transfer rate.

For the primary 29-event R5 cell, uncertainty for paired contrasts was summarized by 5,000 paired replicate bootstrap draws. Replicate indices

Table 3: Frozen H1–H4 support rules in the controlled new-seed experiment.

| ID | Contrast | Primary requirement | Additional frozen requirement |
|---|---|---|---|
| H1 | M2 vs M1 | Mean paired ΔUno-C $\geq +0.02$ in R5 at 29 events | Mean paired direction $\geq 0$ in R5 at both 10 and 40 events |
| H2 | M2 vs M3 | Mean paired ΔUno-C $\geq +0.01$ in R5 at 29 events | ΔIBS $\leq 0$ and Δheld-out risk SD $\leq 0$ |
| H3 | M5 vs M4 | Mean paired ΔUno-C $\geq +0.02$ in R5 at 29 events | Absolute reduction in catastrophic-transfer rate $\geq 0.10$ |
| H4 | M6 vs M4 | Mean paired ΔUno-C $\geq +0.02$ in R5 at 29 events | Absolute reduction in catastrophic-transfer rate $\geq 0.10$ |

were sampled with replacement, and the mean paired difference was recomputed for each draw. The percentile 2.5th and 97.5th percentiles were retained as descriptive 95% intervals. Bootstrap intervals did not change the frozen H1–H4 support rules.

After H1–H4 had been evaluated, a separate arithmetic feasibility audit checked whether each component of the H3/H4 composite rules could have been satisfied given its realized control event rate. This was a post-result diagnostic: it could classify an individual component as structurally unassessable but could not change either frozen machine status or the original threshold.

*S7. Final sign-inversion and regime-shape audit*

The final simulation audit was defined only after H1–H4 were known. It therefore has a weaker status than the controlled H1–H4 experiment. Its interpretation rules were nevertheless written and hashed before reading the new quantities computed specifically for this audit.

For the sign-inversion analysis, the primary 29-event R5 estimand was

$$\Delta C_{\text{sign}} = C(M2) - C(-M1). \tag{7}$$

The result was classified as beyond sign inversion only if the mean difference was at least +0.02 and the lower bound of the paired 95% bootstrap interval was above zero. It was classified as sign-inversion equivalent if the complete 95% interval lay inside $[-0.02, +0.02]$. All other outcomes were classified as inconclusive relative to sign inversion.

Patient-level risk correlation between M2 and M1, cosine similarity between the adapted and source heads, projection of the adapted head onto the source-head direction, and the orthogonal fraction of the adapted head were secondary explanatory quantities. They could not upgrade the prediction-level classification. R0 served as a directional positive control and required both positive median M2–M1 risk correlation and positive median source/adapted-head cosine for geometric interpretation.

The discrete regime-shape audit compared the 29-event transfer gain

$$\Delta C_{M2-B0} = C(M2) - C(M0) \tag{8}$$

in R2 separately with R0, R4, and R5. Because different cells used independent new seeds, independent two-sample bootstrap resampling was used for each other-regime-minus-R2 mean difference. A discrete heterogeneity valley required the lower bound of all three 95% bootstrap intervals to exceed zero. R4 was designated the key anti-low-signal-magnitude comparison because it contains no transportable source component. The rule was explicitly defined over four categorical generator regimes and did not authorize a continuous U-shaped biological claim.

A final M6-prime diagnostic substituted the true binary transportability mask into the already fitted M4 model at prediction time while retaining the M4 residual unchanged. No parameter was refitted. This diagnostic is asymmetric: the retained residual had been optimized jointly with the learned M4 soft gate, not with the substituted truth gate. M6-prime therefore cannot be interpreted as an optimal learned-versus-true-gate comparison and cannot change H4.

The same audit froze a prospective rule for future rate-reduction criteria: future criteria should use a prespecified relative reduction and a prespecified minimum control-rate floor. This future policy was not applied retroactively to H3 or H4.

No additional simulation model branch, A6 reopening, model selection, or reserved human outcome access was permitted after this audit.

*S8. Outcome-free DOG$^2$–TARGET representation*

The real-data expression representation was frozen after the final simulation stop state but before TARGET outcome values were opened.

The primary dog-to-human bridge contained 11,815 one-to-one aligned gene pairs. The exact feature order, DOG$^2$ raw feature, human gene symbol, orthology identifiers, and TARGET expression feature were stored in a frozen alignment artifact. Post-outcome mapping changes were prohibited.

The model representation used all 50 MSigDB Hallmark gene sets [6]. Hallmark membership was frozen before outcome opening. No whole-cohort TARGET Hallmark matrix was calculated.

The raw DOG$^2$ and TARGET matrices were materialized as float64 samples-by-gene matrices in the exact 11,815-gene order. The DOG$^2$ matrix contained 186 samples and the TARGET matrix 88 expression samples. This materialization performed no imputation, variance filtering, scaling, module scoring, PCA, CORAL, outcome-based restriction, or sample deletion.

All TARGET predictive preprocessing was partition-local. For each training partition, gene means and population standard deviations were estimated from training samples only. A gene with training SD $\leq 10^{-12}$ was omitted in that partition. The training parameters were then applied unchanged to the validation or test samples.

For Hallmark $m$ and patient $i$,

$$H_{im} = \frac{1}{|G_m^*|} \sum_{j \in G_m^*} \frac{x_{ij} - \mu_{j,\text{train}}}{\sigma_{j,\text{train}}}, \tag{9}$$

where $G_m^*$ contains mapped genes that remained non-constant in the training partition. At least 10 genes were required for every Hallmark. Hallmark-level means and population SDs were subsequently fitted on the same training partition and applied unchanged to held-out samples.

Inner-CV transformations were fitted separately inside each inner-training fold. After hyperparameter selection, all gene and Hallmark transformations were refitted using the complete outer-training set before the held-out outer fold was transformed.

The full DOG$^2$ source models were permitted to use all 186 source dogs for source-only normalization and source fitting. Source and target domains were standardized independently. Thus, frozen source coefficients or networks were applied to target features represented on the scale defined by the current TARGET training partition; no held-out TARGET observation contributed to this transformation.

*S9. Frozen TARGET model registry*

The complete TARGET model registry was fixed before human outcomes were opened. It contained nine models:

- T0: TARGET-only Hallmark ridge Cox;
- T1: frozen canine Hallmark ridge Cox applied without TARGET outcome fitting;
- T2: frozen canine risk plus a TARGET-fitted residual Cox component;
- N0: frozen canine neural encoder and frozen canine prognostic head;
- N1: frozen canine encoder with a source-initialized, source-centered TARGET prognostic head;
- N2: the N1 target-head procedure with the source-centering penalty removed;
- N3: the original frozen A2 rank-4 residual adapter;
- N4: TARGET-only neural survival model trained from scratch;
- N5: full fine-tuning from the frozen canine neural initialization.

Real-data A3 and hardened A3 were excluded because no valid module-level evolutionary prior had been frozen for TARGET before human outcome access. Neither model could be reconstructed after outcomes were opened.

The canine classical source coefficient vector was materialized before outcome opening using the previously frozen source penalty, without source hyperparameter retuning. The canine neural source network used the exact frozen 50–32–16 architecture and source optimization constants from the simulation implementation.

T0 and T2 used four-fold event-stratified inner cross-validation within each outer-training set. The candidate ridge grid contained 17 logarithmically spaced values from $10^{-4}$ to $10^{4}$. The selected target penalty was determined exclusively from the corresponding outer-training set.

The TARGET neural procedures reused the already-frozen A0/A1/A2/A4 implementation and scientific constants rather than introducing a human-specific architecture search. N2 differed from N1 only by removing the frozen source-head centering penalty. No neural hyperparameter was selected on TARGET.

#### *S10. TARGET endpoint reconciliation and zero-time amendment*

The outcome-blind TARGET expression roster contained 88 samples. After the primary OS endpoint was opened, two samples lacked the information required by the already-existing complete-case OS policy. The resulting primary analysis population contained 86 cases, 29 observed events, and 57 censored observations. This population reproduced the pre-existing complete-case TARGET OS population rather than defining a new exclusion rule.

No additional clinical variable was read to remove a case, and no sample was excluded according to expression, model behavior, or later performance.

One of the 86 cases, TARGET-40-PAKUZU, was censored at an observed OS time of exactly zero. The value was retained without imputation, epsilon shifting, or sample exclusion.

Before fitting any real TARGET model, the installed survival stack was audited using synthetic zero-time cases and the exact 100 frozen 20-repeat-by-5-fold TARGET split configurations. The survival-object constructor supported the censored zero-time value in all 100 folds, and finite Uno-C support was verified in all 100 folds. The zero-time case occurred in the training set of 80 folds and the test set of 20 folds. The existing IBS support/nonassessability policy was audited separately and was not changed.

The pre-outcome runner contained two assumptions that no longer matched the reconciled endpoint: it expected 88 complete outcomes and rejected all times less than or equal to zero. Exactly three technical substitutions were therefore authorized:

1. expected TARGET endpoint count: $88 \rightarrow 86$;
2. expected TARGET expression shape: $(88, 11815) \rightarrow (86, 11815)$;
3. invalid-time guard: $\text{time} \leq 0 \rightarrow \text{time} < 0$.

No model, hyperparameter, metric, resampling, bootstrap, endpoint, interpretation branch, or other runner logic was permitted to change.

The first technical execution attempt failed before model fitting because Windows text-mode newline translation changed the raw runner bytes. The next wrapper changed patch construction to raw-byte mode while preserving the source newline convention and required reverse patching to reproduce the pre-outcome runner byte-for-byte.

A subsequent preflight failure arose because a whole-file SHA-256 lock on a rerunnable zero-time audit JSON included runtime metadata. The final execution wrapper did not replace that historical hash with a newly observed value. Instead, it required the run-local summary to cryptographically bind the current audit and independently replayed the exact 86/29/57 endpoint semantics, zero-time support, child-artifact hashes, and the three authorized substitutions. These corrections occurred before real TARGET model fitting and introduced no scientific change.

*S11. TARGET repeated cross-validation and metrics*

All nine TARGET models used identical event-stratified outer folds. The evaluation used five outer folds repeated 20 times. Repeat $r$ used the fixed seed

$$20260830 + r. \tag{10}$$

An individual patient therefore appeared exactly once as a held-out observation within each repeat, but the resulting 20 predictions were not treated as 20 independent biological observations.

Outer-test samples were used only for transformation under parameters fitted on the corresponding outer-training set and for prediction. No outer-test outcome entered hyperparameter selection, preprocessing, or refitting.

The primary TARGET metric was Uno-C [2]. The human evaluation used the 90th percentile of outer-training event times as its time-support horizon. This value had been frozen separately from the 80th-percentile simulation rule.

Secondary TARGET metrics were IBS, held-out risk-score population SD, and the 99th-to-1st percentile risk-score range. For IBS, a Breslow baseline cumulative hazard was estimated from the original outer-training outcomes for the fixed model risk score. This recalibration did not refit the model's risk coefficient, slope, intercept, or ranking.

Within a repeat, the five outer-fold metric values were combined by sample-count weighting. The final model estimate was the arithmetic mean of valid repeat-level estimates. A minimum of 16 valid repeats out of 20 was frozen before outcome opening. If fewer than 16 repeats supported a required metric, that metric was classified as not assessable.

Paired model contrasts were calculated within the identical fold and repeat before applying the same fold-to-repeat and repeat-to-final aggregation.

*S12. Patient-clustered TARGET bootstrap*

TARGET uncertainty was calculated from fixed repeated-CV predictions rather than by refitting the entire model-development process.

The bootstrap sampling unit was the unique TARGET patient/case key. Patients were stratified by OS event status and sampled with replacement within the event strata. A sampled patient's multiplicity was propagated identically across all models and across all 20 held-out appearances of that patient.

For a patient selected more than once in a bootstrap draw, the held-out outcome and risk row was repeated literally according to the sampled multiplicity when the fold metric was recomputed. Original outer-training survival distributions were retained. Models were not refitted, preprocessing was not repeated, and outer splits were not regenerated.

For each bootstrap draw, fold metrics were recomputed from the resampled held-out rows and then aggregated using the same sample-count-weighted fold-to-repeat and arithmetic repeat-to-final rules as the point estimate. Paired contrasts were calculated from the same draw and same patient multiplicities for both models.

The deterministic bootstrap stream continued until 5,000 valid draws were obtained or 25,000 total attempts had been made. Percentile 95% intervals were the 2.5th and 97.5th percentiles of the valid bootstrap distribution. If 5,000 valid draws could not be obtained for a branch-defining quantity, that comparison was not assessable and the descriptive branch was assigned T-D.

These intervals quantify sampling uncertainty conditional on the frozen repeated-CV predictions. They are not confidence intervals for the complete model-development procedure.

*S13. Frozen TARGET interpretation states*

TARGET interpretation used the neural N0/N1 mechanism arm with T0 as the target-only reference. The following point-estimate states were frozen before outcome opening.

T-A, the concordant-source pattern, required

$$C(N0) > 0.50, \qquad \Delta C(N1 - N0) < +0.02, \\ \Delta C(N0 - T0) > -0.02. \tag{11}$$

T-B, the reversal-like adaptation pattern, required

$$C(N0) < 0.50 \quad \text{and} \quad \Delta C(N1 - N0) \geq +0.02. \tag{12}$$

If neither T-A nor T-B applied, T-C required

$$\Delta C(N0 - T0) < +0.02 \quad \text{and} \quad \Delta C(N1 - T0) < +0.02. \tag{13}$$

T-D represented an inconclusive or unresolved pattern. T-D was also assigned mechanically if a condition required by the point-estimate branch was uncertain under the frozen bootstrap rules. Specifically, orientation was unresolved if the 95% interval for $C(N0)$ included 0.50, and a materiality condition was unresolved if the corresponding paired $\Delta C$ interval contained both $-0.02$ and $+0.02$.

The branch assignment was mechanical and did not allow subjective override. T-D was a valid expected result rather than a failure of the analysis.

Additional reporting flags described direct classical or neural source benefit, positive concordance after target-head retargeting, and disagreement in orientation between classical and neural outcome-zero-shot source scores. These flags could not change the branch.

No multiplicity correction was used across the nine TARGET models because this entire analysis was designated descriptive and non-confirmatory. No individual model-wise significance result could select a model or change the prior simulation conclusion.

*S14. Secondary outcome-blind biological-context analysis*

The biological-context analysis was frozen after TARGET outcomes had technically been opened but before completed TARGET model results were inspected. Its evidentiary status is therefore secondary and post-opening, but the analysis itself is outcome-blind.

The primary landscape contains all 50 frozen Hallmark gene sets. It uses all 186 DOG$^2$ expression samples and all 88 TARGET expression samples and does not intersect TARGET with survival completeness. Whole-cohort use is permitted here because the analysis does not fit a survival predictor, calculate held-out prediction performance, or use survival outcomes.

For each gene set, a Pearson gene–gene correlation matrix is calculated separately in DOG$^2$ and TARGET. Genes are placed in the same sorted human gene-symbol order in both cohorts, and the strict upper triangle is extracted from each matrix. The primary structural statistic is

$$S_m = \rho_{\text{Spearman}}\left(\mathbf{e}_{D,m}, \mathbf{e}_{H,m}\right), \tag{14}$$

where $\mathbf{e}_{D,m}$ and $\mathbf{e}_{H,m}$ denote the DOG$^2$ and TARGET edge vectors for gene set $m$. Pearson edge concordance and non-zero edge-sign agreement are secondary descriptive statistics.

Each gene set is compared with 1,000 exact-size random panels drawn from the 11,815-gene aligned universe. Genes belonging to the tested set are excluded from its control pool.

Random controls are matched jointly for outcome-blind expression level and variance in both cohorts. Four gene-level quantities are calculated: DOG$^2$ median-expression rank, DOG$^2$ variance rank, TARGET median-expression rank, and TARGET variance rank. Average ranks are rescaled as

$$q = \frac{\text{rank} - 1}{N_{\text{genes}} - 1} \tag{15}$$

and divided into the intervals $[0, 1/3)$, $[1/3, 2/3)$, and $[2/3, 1]$. The Cartesian product gives $3^4 = 81$ joint strata.

For an observed gene set, every matched random panel must reproduce exactly its full 81-stratum count vector. Sampling is without replacement within a panel and independent across panels. No nearest-neighbor, unmatched, or relaxed fallback is permitted; inability to construct the required matched controls causes the analysis to fail closed.

For statistic $S_m$, the matched-control percentile is

$$P_m = \frac{1 + \sum_{b=1}^{1000} I(S_{m,b}^{\text{random}} \le S_m)}{1001}. \tag{16}$$

This quantity is a descriptive percentile and is not a $p$-value. Random median, 5th percentile, and 95th percentile are also retained.

The 50 Hallmark sets overlap in gene membership. All 1,225 Hallmark pairwise overlaps are therefore audited, and the 50 modules are not treated as independent observations. No module-level regression or correlation $p$-value, and no Bonferroni or Benjamini–Hochberg correction treating the modules as an independent $n = 50$, is authorized.

Four molecular programs (M34, M40, M11, and M24) from a separately conducted outcome-blind cross-cohort preservation analysis were carried forward as frozen coherence anchors. Their memberships and preservation labels were fixed before the present biological-context analysis and were not modified using TARGET outcomes or the current structural-concordance results.

M34 and M40 entered with frozen labels of strong external canine representation preservation, whereas M11 and M24 entered with frozen labels of no clear external preservation. The anchors were retained regardless of whether the current structural statistic agreed with those prior labels and were not used for model selection.

The anchor comparison is a coherence check rather than an independent validation test. Disagreement cannot trigger removal of an anchor, alteration of the structural statistic, or selection of a replacement anchor.

The structural concordance statistic is also not independent of the type of expression organization that a cross-species representation can exploit. Accordingly, the analysis characterizes the biological/domain-shift setting and cannot be used to claim that structural concordance independently predicts TARGET transfer utility.

A module-level diagnostic derived from the completed TARGET model output is permitted only if the frozen model output already admits an exact Hallmark decomposition without approximation, parameter re-estimation, model refitting, new prior construction, or a new gate. Forced Hallmark attribution of hidden N1/N2 representations, post-hoc Hallmark attribution of the N3 low-rank adapter, and construction of a new real-data A3 model are prohibited. If the decomposition condition is not met, the module-level TARGET diagnostic is omitted.

A null or mixed biological result remains reportable. It cannot trigger reoptimization of the statistic, random controls, or anchors. If the biological result does not justify a full main-text figure, the reserved Figure 5 position may return to the already completed gating analysis without changing the evidentiary status of the biological analysis.

*S15. Software, execution devices, and reproducibility*

The analysis was implemented in Python using NumPy, pandas, PyTorch [7], scikit-survival [3], and scikit-learn where required. Exact package and platform versions used for final analyses are reported in Supplementary Table S7.

The large 21,600-replicate simulation model matrix and the 6,000-replicate controlled model matrix used GPU execution where specified by their frozen implementation contracts. Metric aggregation and diagnostic analyses that did not benefit materially from GPU execution were run on CPU.

The final TARGET evaluation retained the audited CPU execution path because the pre-outcome runner and zero-time survival-stack validation had been frozen on that path. Execution device was not used as a model-selection variable.

Simulation recipes and model outputs were written to restart-safe, scenario- or cell-specific shards with SHA-256 manifests. Downstream scripts verified both the expected prior frozen contract hash and the individual artifact hash before reading a scientific result.

The TARGET analysis used the same principle. The pre-outcome source artifacts, representation contract, clinical-schema contract, model implementation inventory, executable runner, endpoint reconciliation, zero-time audit, and technical runner amendment all have separate immutable provenance records.

The biological-context contract similarly records the exact Hallmark and anchor membership hashes, aligned matrix hashes, random-control rules, and the filesystem chronology demonstrating that the contract was frozen before a completed TARGET result was available.

No original frozen artifact is deleted from the reproducibility record when a technical amendment is required. Earlier failed technical attempts are retained with their failure stage and whether any scientific fitting occurred before the failure. The final released repository will include the scripts, configuration files, frozen registries, seed manifests, major derived non-identifying artifacts, and hash ledger required to reconstruct the reported analysis.

*S16. Post-HOLD manuscript-strengthening sensitivity analyses*

After the original simulation HOLD and all model-development stop rules had already been fixed, a separate strengthening contract specified a bounded set of additional summaries intended to address likely reviewer questions without reopening model selection. These analyses were explicitly post-HOLD and could not make a nonselectable architecture selectable, change the frozen $-0.02/-0.05$ harm definitions, or alter any TARGET interpretation state.

The first analysis exported the complete seven-model frozen safety summary from the already computed scenario-level metrics. No survival metric was recomputed. The second analysis compared the original 180-scenario equal-weight estimand with two fixed alternatives: the 36 pre-stress core scenarios with equal weight, and a regime-balanced estimand assigning each

**Supplementary Table S9. Descriptive all-model frozen simulation safety summary.**

| Model | Selectable | Mean Uno-C | Mean $\Delta C$ vs B0 | NT rate | Catastrophic rate |
|---|---|---|---|---|---|
| B0 | No | 0.555554 | 0.000000 | 0.000000 | 0.000000 |
| B4 | No | 0.544989 | −0.010566 | 0.305528 | 0.099444 |
| A0 | No | 0.555351 | −0.000203 | 0.009000 | 0.000000 |
| A1 | No | 0.572198 | +0.016644 | 0.286778 | 0.160556 |
| A2 | Yes | 0.569809 | +0.014254 | 0.215417 | 0.071972 |
| A3 | Yes | 0.530067 | −0.025487 | 0.486333 | 0.326889 |
| A4 | No | 0.519107 | −0.036447 | 0.511722 | 0.407833 |

NT uses $\Delta C \leq -0.02$ and the overall catastrophic rate uses $\Delta C \leq -0.05$, both with equal scenario weighting across the frozen 180-scenario benchmark. Eligibility is the original frozen architecture-selection eligibility and is not changed by this descriptive table.

of R0–R5 weight 1/6 after within-regime averaging. Because retained per-replicate metric arrays existed, Monte Carlo uncertainty was summarized with 4,000 deterministic bootstrap draws that resampled replicates within each fixed scenario. Scenarios were never resampled and the intervals therefore quantify finite-replicate Monte Carlo uncertainty for the fixed benchmark rather than uncertainty over a hypothetical scenario superpopulation.

A third sensitivity evaluated neighboring definitions of negative and catastrophic transfer. Negative-transfer severity was evaluated at $\Delta C \leq -0.01, -0.02, -0.03$ and catastrophic-transfer severity at $\Delta C \leq -0.04, -0.05, -0.06$. The original $-0.02$ and $-0.05$ definitions remained the only frozen primary definitions and were required to reproduce the existing benchmark rates exactly.

Finally, a post-opening no-covariate TARGET reference was calculated to contextualize IBS. The exact frozen 20-repeat-by-5-fold partitions were reused. For each outer fold, a Kaplan–Meier survival curve was estimated from that fold's outer-training outcomes only and assigned unchanged to every held-out patient. The existing 04c IBS time-grid, censoring/IPCW implementation, and sample-count-weighted fold-to-repeat/arithmetic repeat-to-final aggregation were reused exactly. Manual product-limit and scikit-survival Kaplan–Meier implementations agreed to machine precision in all folds. This reference was descriptive only and could not be used for model selection or T-A–T-D branch assignment.

**Supplementary Table S10. Aggregation sensitivity for A2 and A3.**

| Estimand | Model | Mean $\Delta C$ [MC 95%] | NT rate [MC 95%] | Catastrophic rate [MC 95%] |
|---|---|---|---|---|
| Frozen original | A2 | 0.014254 [0.013653, 0.014835] | 0.215417 [0.209833, 0.221083] | 0.071972 [0.068472, 0.075528] |
| Frozen original | A3 | −0.025487 [−0.025956, −0.025022] | 0.486333 [0.482306, 0.490278] | 0.326889 [0.323056, 0.330834] |
| Core only | A2 | 0.019287 [0.018193, 0.020372] | 0.209583 [0.200417, 0.218611] | 0.085694 [0.079444, 0.092083] |
| Core only | A3 | 0.003248 [0.002518, 0.003946] | 0.324722 [0.317639, 0.331944] | 0.153056 [0.147222, 0.158889] |
| Regime balanced | A2 | 0.015577 [0.014645, 0.016453] | 0.219263 [0.211601, 0.227083] | 0.085374 [0.080000, 0.090962] |
| Regime balanced | A3 | −0.001132 [−0.001767, −0.000542] | 0.343718 [0.337308, 0.350342] | 0.173547 [0.169134, 0.178120] |

MC intervals resample retained replicates within fixed scenarios; scenarios are not resampled. The frozen-original estimand remains primary.

[7] A. Paszke, S. Gross, F. Massa, A. Lerer, J. Bradbury, G. Chanan, T. Killeen, Z. Lin, N. Gimelshein, L. Antiga, A. Desmaison, A. Köpf, E. Yang, Z. DeVito, M. Raison, A. Tejani, S. Chilamkurthy, B. Steiner,

**Supplementary Table S11. Neighboring harm-definition sensitivity.**

| Definition family | Model | Severity threshold | Equal-scenario rate | Change vs frozen definition |
|---|---|---|---|---|
| Negative transfer | A2 | −0.01 | 0.289611 | +0.074194 |
| Negative transfer | A2 | −0.02 | 0.215417 | 0.000000 |
| Negative transfer | A2 | −0.03 | 0.153389 | −0.062028 |
| Negative transfer | A3 | −0.01 | 0.546778 | +0.060444 |
| Negative transfer | A3 | −0.02 | 0.486333 | 0.000000 |
| Negative transfer | A3 | −0.03 | 0.429250 | −0.057083 |
| Catastrophic transfer | A2 | −0.04 | 0.106972 | +0.035000 |
| Catastrophic transfer | A2 | −0.05 | 0.071972 | 0.000000 |
| Catastrophic transfer | A2 | −0.06 | 0.047306 | −0.024667 |
| Catastrophic transfer | A3 | −0.04 | 0.375889 | +0.049000 |
| Catastrophic transfer | A3 | −0.05 | 0.326889 | 0.000000 |
| Catastrophic transfer | A3 | −0.06 | 0.281833 | −0.045056 |

The frozen −0.02 negative-transfer and −0.05 catastrophic-transfer definitions remain the only primary definitions. Neighboring rows are post-HOLD sensitivity summaries only.

L. Fang, J. Bai, S. Chintala, PyTorch: An imperative style, high-performance deep learning library, in: Advances in Neural Information Processing Systems, Vol. 32, 2019.

**Supplementary Table S12. TARGET model IBS relative to a fold-local no-covariate Kaplan–Meier reference.**

| Model | Existing IBS | KM-null IBS | Model − KM-null |
|---|---|---|---|
| T0 | 0.209292 | 0.165645 | +0.043647 |
| T1 | 0.159032 | 0.165645 | −0.006612 |
| T2 | 0.203835 | 0.165645 | +0.038190 |
| N0 | 0.236592 | 0.165645 | +0.070947 |
| N1 | 0.169071 | 0.165645 | +0.003426 |
| N2 | 0.169098 | 0.165645 | +0.003453 |
| N3 | 0.204556 | 0.165645 | +0.038911 |
| N4 | 0.238839 | 0.165645 | +0.073195 |
| N5 | 0.234114 | 0.165645 | +0.068469 |

The KM reference is fitted separately in each frozen outer-training fold and assigns the same training-derived survival curve to every corresponding outer-test patient. Lower IBS is better. Comparisons are descriptive only.

**Supplementary Table S13. Chronology and provenance of pre-existing biological anchors.**

| Program | Aligned genes | Pre-existing preservation label | Provenance and chronology evidence |
|---|---|---|---|
| M34 | 162 | Strong external canine representation preservation | Membership and label fixed in an earlier outcome-blind analysis by the same authors |
| M40 | 111 | Strong external canine representation preservation | Membership and label fixed in an earlier outcome-blind analysis by the same authors |
| M11 | 7 | No clear external canine representation preservation | Membership and label fixed in an earlier outcome-blind analysis by the same authors |
| M24 | 7 | No clear external canine representation preservation | Membership and label fixed in an earlier outcome-blind analysis by the same authors |

The four programs were not selected using the current Hallmark landscape or TARGET survival results. Their memberships were bound to the earlier frozen gene-weight artifact (SHA-256 prefix `4f065aa4c4edf117`), and their pre-existing labels were bound to the corresponding frozen manifest (prefix `fcd34b985ea99986`). Before the present structural analysis was executed, the biological-context contract (SHA-256 `405d6d6e6c8f2a7dfeb85d32c7ac045543e9e1ea8b61458e74d166367b4370ea`) verified that the completed TARGET model-result summary was absent at both the initial gate and immediately before contract write, and recorded a contemporaneous filesystem snapshot. The subsequent outcome-blind execution replayed the same anchor definitions without reading the completed TARGET scientific results. Exact filesystem timestamps, complete artifact hashes, and the full hash chain are retained in the released provenance ledger.
